# AFM-based force spectroscopy unravels the stepwise-formation of a DNA transposition complex driving multi-drug resistance dissemination

**DOI:** 10.1101/2022.07.18.500257

**Authors:** Maricruz Fernandez, Alexander V. Shkumatov, Yun Liu, Claire Stulemeijer, Sylvie Derclaye, Rouslan G. Efremov, Bernard Hallet, David Alsteens

## Abstract

Transposon Tn4430 belongs to a widespread family of bacterial transposons, the Tn3 family, which plays a prevalent role in the dissemination of antibiotic resistance among pathogens. So far, the molecular mechanisms underlying the replicative transposition of these elements are still poorly understood. Here, we use force-distance curve-based atomic force microscopy to probe the binding of the TnpA transposase of Tn4430 to DNA molecules containing one or two transposon ends and to extract the thermodynamic and kinetic parameters of transposition complex assembly. Comparing wild-type TnpA with previously isolated deregulated TnpA mutants supports a stepwise pathway for transposition complex formation and activation during which TnpA first binds to a single transposon end and then undergoes a structural transition that enables it to bind the second end co-operatively and to become activated for transposition catalysis. Our study thus provides an unprecedented approach to probe the dynamic of a complex DNA processing machinery at the single-particle level.

## INTRODUCTION

Transposable elements or ‘transposons’ are widely diverse genetic elements that can propagate from one location to another within the genome of their host (1). This movement is mediated by ‘transposases’, specialized recombination enzymes to cut and paste DNA molecules (1, 2). Transposase genes represent the most abundant and ubiquitous annotated genes in genomes, which reflects the evolutionary impact of transposable elements throughout the living world, being responsible for deleterious mutations and diseases, but also the creation of new genes, the establishment of complex regulatory networks, and the exchange of useful phenotypic traits through horizontal gene transfer (3–7). As such, transposable elements play a prominent role in the emergence of new pathogens and the dissemination of antimicrobial resistance among bacteria (8).

In particular, transposons of the wide-spread Tn*3*-family pose a permanent threat to human health, being continuously involved in the dispersal of new resistances, including against last-chance antibiotics such as carbapenems and colistin (8–12). The effectiveness of these transposons relies on their ability to copy themselves and their passenger genes through replicative transposition every time they move (8). The transposases (TnpA) of Tn3-family transposons form a distinct subgroup of the structurally diverse ‘DDE/D’ transposase/integrase superfamily that cleave and reseal DNA molecules using a triad of acidic residues (DDE or DDD) imbedded in a conserved RNaseH-like catalytic domain (13–16). In the case of the Tn3-family, the strand transfer intermediate resulting from the joining of the transposon 3’-ends to the target DNA is processed by the host replication machinery to generate two new copies of the transposon. However, despite the biological and societal impact of Tn3-family transposons, the molecular mechanism whereby TnpA interacts with the DNA partners to assemble the active transposition complex (or ‘transpososome’) and catalyze the reaction remains poorly understood (9).

Our recent work on the Tn3-family member Tn4430 has opened up new avenues toward the understanding of this mechanism (17, 18). Biochemical characterization of the early stages of transposition supported a unique and asymmetric scenario for transposase assembly and activation (Fig. 1*A*). This scenario was established by comparing the wild-type TnpA of Tn4430 (TnpA^WT^) with deregulated TnpA mutants (TnpA^S911R^ and the triple mutants Tnp^AW24R/A174V/E740G^, hereafter referred to as TnpA^3X^) that had previously been isolated for their impairment in target immunity, a yet poorly understood TnpA-dependent mechanism whereby Tn3-family transposons avoid inserting multiple times into the same target (13, 19). TnpA^S911R^ and TnpA^3X^ displayed promiscuous target selectivity *in vivo*, promoting transposition into an immune target at a higher frequency than TnpA^WT^, and proved to be hyperactive in different biochemical assays *in vitro*.

**Fig 1.**
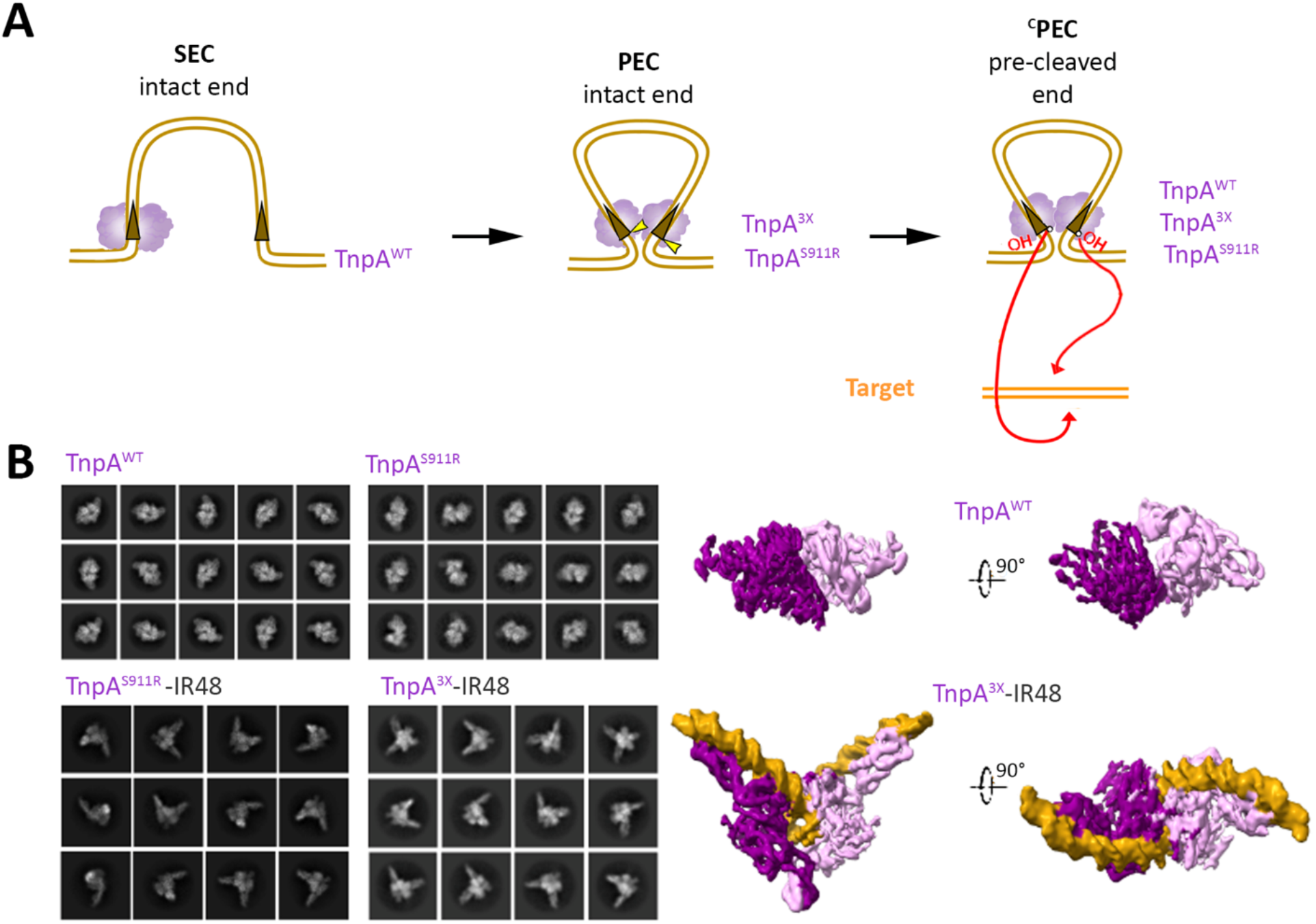
Probing Tn4430 transposition by AFM-based force spectroscopy. (A) Model for Tn4430 transposition pathway. TnpA (purple) primarily binds to the transposon IR end (brown triangle) to assemble a single-end complex (SEC). TnpA undergoes an activating conformational change to assemble a paired-end complex (PEC) where it is competent for DNA cleavage (yellow arrowheads). Cleavage exposes a 3’OH group at both transposon ends, which are then transferred (red arrows) to staggered positions of target DNA. In vitro, TnpA^WT^ can only form PEC on pre-cleaved ends (^c^PEC), while the hyperactive TnpA^S911R^ and TnpA^3X^ mutants spontaneously assemble PEC and ^c^PEC on both intact and pre-cleaved ends. (B) 2D class averages for TnpA^WT^, TnpA^S911R^ and TnpA^S911R^-IR48, TnpA^3X^-IR48 (left pane). Medium resolution models for TnpA^WT^ (4.4 Å) and TnpA^3X^-IR48 (6.8 Å) showing dimeric structure of TnpA (right panel). Monomers are colored in light and dark purple, IR48 substrate is colored in dark yellow.

TnpA^WT^ was shown to specifically bind to a DNA fragment carrying the terminal inverted repeat (IR) of Tn4430 to form a single-end complex (SEC), whereas TnpA^S911R^ and TnpA^3X^ spontaneously and cooperatively assembled a paired-end complex (PEC) containing two IR ends. Formation of the PEC correlated with activation of DNA cleavage at the transposon ends by TnpA (17). TnpA^WT^ can only form the PEC on precleaved ends *i*.*e*. when the initial step of transposition is bypassed, whereas the hyperactive mutants TnpA^S911R^ and TnpA^3X^ can overcome the activation barrier for PEC assembly on both cleaved and un-cleaved (intact) ends (17). Together, the data suggest that transposition is normally controlled at a critical stage of transposition complex (or ‘transpososome’) assembly, before DNA cleavage, and that activation of TnpA involves a conformational change that promotes the formation of a stable synaptic complex (PEC) to trigger catalysis and coordinate DNA cleavage and strand transfer reactions at both transposon ends (Fig. 1*A*). However, due to the lack of direct evidence, the dynamic of complex formation and the requirement for a structural transition toward the active conformer remained hypothetical.

Here, we applied force-distance (FD)-based atomic force microscopy (AFM) (20) to analyze the binding of TnpA and in particular, to compare the binding of TnpA^WT^ with the previously characterized TnpA^S911R^ and TnpA^3X^ deregulated variants. To this end, we engineered DNA molecules containing one or two Tn4430 IR ends, to decipher transposition complex assembly at the single particle level. We extracted the kinetics and thermodynamics of TnpA-DNA interactions and probed conformational changes taking place during TnpA-DNA complex formation. The findings provide new quantitative insights on Tn4430 transposition complex formation with a resolution that could hardly be reached using conventional biochemical approaches, thus constituting a pioneering study in the field of protein-DNA transactions.

## MATERIALS AND METHODS

### Bacterial strains, plasmids, and oligonucleotides

The strains, plasmids, and oligonucleotides used in this study are listed in *SI Appendix*, Table S3, and S4, respectively.

### TnpA expression and purification

TnpA proteins fused to a cMyc-His_6_ epitope tag at their C terminus were expressed in *E. coli* TOP10 cells under the control of the p*Ara* promoter (*SI Appendix*, Table S3). TnpA derivatives were purified using the thermal shift-induction procedure to increase the yield of soluble proteins followed by nickel column chromatography as described(13, 18). After purification, proteins were washed with the lysis buffer [50 mM Tris·HCl (pH 8.0), 500 mM NaCl and 20% (vol/vol) glycerol], and concentrated using an Amicon Ultra-0.5 mL 50 K filter (Merck).

### Small-angle X-ray scattering (SAXS) data collection and analysis

Small-angleX-ray scattering (SAXS) data were collected on the SWING beamline (21) at Soleil synchrotron (Gif-sur-Yvette, France). The TnpA^WT^ containing fractions [50 mM HEPES (pH 7.9), 1 M NaCl, 10% glycerol, ∼400mM imidazole] after elution from HisTrap column were concentrated to ∼3.6 mg/ml and ∼50ul of the sample was loaded onto a BioSEC3-300 (Agilent) column equilibrated in SEC buffer [50 mM HEPES (pH 7.9), 200 mM NaCl, 100 mM L-Arg HCL]. During the elution, 600 scattering measurements were taken with 1*s* time frames. Data reduction, frame averaging, buffer subtraction, and initial analysis were performed using FOXTROT (21) and DATASW (22) programs. Further interpretation of the SAXS data involved representation using the dimensionless Kratky plot (23) and calculation of the pair-distance distribution function using the ATSAS program GNOM (24). The excluded volume was estimated using DAMMIN (25) in P1 and P(r) function. The data collection parameters and calculated invariants are summarized in *SI Appendix*, Table S1.

### Cryo-EM grids preparation, imaging and processing

Freshly gel filtered sample (TnpA^S911R^ or TnpA^WT^) was used for grid preparation or in case of TnpA^S911R^ and TnpA^3X^ were mixed with a 4-10-fold excess of IR48 and subjected to a second gel filtration to remove unbound substrate. The grids were prepared by applying 5µl of the sample on glow-discharged Quantifoil R2/1 mesh 300 or UltrAufoil R1.2/1.3 mesh 300 grids at concentrations of ∼0.1 and ∼0.6-1 mg/ml, respectively. After 60s incubation, the excess of the sample was removed by blotting from one side for 2.3s at 80-90% relative humidity and plunge frozen in liquid ethane using a Cryoplunge 3 System (Gatan).

Micrographs were collected at 300 kV on a CRYO ARM 300 (JEOL) electron microscope at a nominal magnification of 60,000 and corresponding pixel size of approximately 0.76 Å. The images were recorded using a K3 detector (Gatan) operating in correlative-double sampling (CDS) mode. The microscope illumination conditions were set to spot size 6, alpha 1, and the aperture diameters for the condenser and objective apertures were 100 and 150 µm, respectively. The energy filter slit was centered on the zero-loss peak with a slit width set to 20 eV. Coma-corrected data acquisition (26) was used to acquire between 6 and 25 micrographs per stage position using SerialEM v3.0.8 (27). Each micrograph was recorded as a movie of 61 frames over a 3 s exposure time and at a dose rate of 11 e^-^pixel^-1^s^-1^ (corresponding to a dose rate per frame of 0.6 e^-^Å^-2^) and total exposure dose of approximately 60 e^-^Å^-2^.

Initial data processing was performed on-the-fly using RELION_IT (28). Dose-fractionated movies were subjected to motion correction and dose weighting using the MotionCorr2 (29). The dose-weighted aligned images were used for CTF estimation using CTFFIND-4 (30). An in-house script was used to plot the calculated parameters, visualize the results, and select the micrographs for further processing (31). The aligned and dose-weighted images were imported into cryoSPARC v3.1.0 (32), and CTF was calculated using Patch CTF. Particle selection was performed using a blob or template-based picker followed by several rounds of 2D classification. The 2D classification, *ab initio* reconstruction and 3D refinement were performed using cryoSPARC. The data collection parameters are summarized in *SI Appendix*, Table S2.

### Functionalization of AFM tips

OMCL-TR400PB-1 cantilevers (Olympus) were used to probe the interaction between TnpA^WT^, TnpA^3X,^ and TnpA^S911R^ and the DNA molecule containing one or two transposon ends. To functionalize the AFM tips, the cantilevers were rinsed with ethanol, dried under a gentle stream of nitrogen, cleaned for 15 min by UV and ozone treatment (Jetlight), and incubated overnight in a solution of 0.1 mM HS-(CH_2_)_11_-EG_3_-OH and 0.1 mM HS-(CH_2_)_11_-EG_3_-NTA (99:1). Then, they were rinsed with ethanol, dried with nitrogen, and immersed for 30 min in 50mM NiCl_2_ before using (33).

The tips were functionalized with the TnpA proteins through their (His)6 purification tag. The cantilevers were placed on Parafilm (Bemis NA), incubated with 50 μl of a TnpA protein solution (0.1 mg/ml) for 1 h, and washed three times with DNA binding buffer [50mM HEPES and 100mM NaCl].

### Preparation of DNA-coated model surfaces

DNA molecules containing one or two transposon ends were immobilized using click chemistry as described (34). Briefly, gold-coated surfaces were rinsed with ethanol, dried under a gentle stream of nitrogen, cleaned for 15 min by UV and ozone treatment (Jetlight), and incubated overnight in a solution of 0.1 mM HS-(CH_2_)_11_-EG_3_-OH and 0.1 mM HS-(CH_2_)_11_-EG_5_-N_3_ (95:5). Then, they were rinsed with ethanol and dried with nitrogen. Immediately afterward, a 30 μl aliquot containing 30 µM of the one-end or two-end substrate (with a terminal or internal alkyne group, respectively), 1 M ascorbic acid, and 0.3 M CuSO_4_ was deposited on the surfaces and left to incubate overnight in the dark at 4 °C. Finally, the surfaces were washed three times with DNA binding buffer.

### Force-distance- (FD-) based AFM on model surfaces

FD-based AFM on model surfaces was performed in DNA binding buffer at room temperature, with functionalized OMCL probes (Olympus, spring constants from 0.027 to 0.053 Nm^−1^ calculated using thermal tune) as described above (35). A Force-Robot 300 (Bruker, Germany) operated in the force-volume (contact) mode was used. Areas of 5 × 5 μm were scanned, the approach and retraction speeds were 1 μm/s, the hold time was 200 ms, the ramp size was set to 300 nm and the setpoint force at 250 pN, with a resolution of 32 × 32 pixels.

Dynamic force spectroscopy (DFS) analysis was performed using a constant approach speed of 1 μm/s and variable retraction speeds (0.1, 0.2, 1, 2 and 10 μm s^−1^), and kinetic on-rate was estimated by measuring the binding probability (BP; i.e., the fraction of curves showing binding events) at different hold times (*i*.*e*., the time the tip is in contact with the surface; 100, 200, 400, 600, 800, 1000 and 1200 ms). Force curves were analyzed on the data processing software from JPK Data processing (version 6.1.149). To identify peaks corresponding to adhesion events between proteins linked to the PEG spacer and the DNA molecule of the model surface, the retraction curve before bond rupture was fitted with the worm-like chain model for polymer extension (36).

Origin software (OriginLab) was used to display the results in DFS plots, to fit histograms of rupture force distributions for distinct loading rate ranges, and to apply various force spectroscopy models (37–39).

For kinetic on-rate analysis, BP was determined at a certain hold time (t). Those data were fitted and *K*_D_ calculated described (40, 41). In brief, the relationship between interaction time (τ) and BP is described by the following equation:

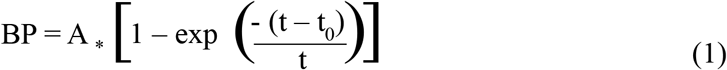

where *A* is the maximum BP and *t*_0_ is the lag time. Origin software was used to fit the data and extract τ. In the next step, *k*_on_ was calculated with the following equation, where *r*_eff_ is the radius of the sphere, *n*_b_ the number of binding partners, and *N*_A_ the Avogadro constant:

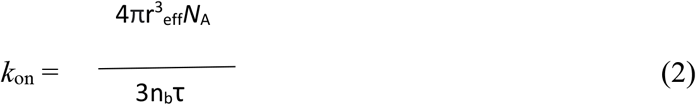

The effective volume *V*_eff_ [(4*πr*_eff_^3^)/3] represents the volume in which the interaction can take place.

### Competition-binding assays

To monitor the effect of a second transposon end (IR end-2 or blocking DNA; see *SI Appendix*, Table S4) on TnpA-one end interaction, BP was measured before and after tip incubation with 12.5, 25, and 50 μM of the blocking DNA. A first map (5 × 5 μm) was recorded as described above (see previous section, FD-based AFM on model surfaces). Then, the appropriate concentration of the second transposon end was injected, and a new map was recorded.

## RESULTS

### Dimeric structure of the free and DNA bound TnpA

PEC formation implies that TnpA has two DNA binding interfaces to complex both transposon ends specifically. This was difficult to reconcile with the finding that the protein-DNA ratio is the same in the SEC and PEC and our previous estimations based on size exclusion chromatography (SEC) and dynamic light scattering (DLS) analysis suggesting that purified TnpA was a monomer in solution (17). However, the data were somewhat ambiguous supporting an ‘extended’ configuration of the protein with an apparent size between that of a monomer and a dimer (17). Since it was necessary to ascertain the binding determinants of TnpA, we have decided to revisit the data by measuring the structural properties of TnpA with greater accuracy using small-angle X-ray scattering (SAXS) coupled to SEC (SEC-SAXS) (*SI Appendix*, Fig. S1 and Table S1). The SEC-SAXS profile of TnpA^WT^ shows a major symmetric peak with a constant radius of gyration (*Rg*) across the elution profile, indicating the absence of interparticle interactions (*SI Appendix*, Fig. S1*A-C*). Further analysis of SAXS data yielded *R*_*g*_ (from Guinier plot) and *D*_*max*_ values of 46 ± 5 and 160 ± 10 Å, respectively, whereas the molecular mass estimation was ∼270-300 kDa (*SI Appendix*, Table S1), which is closer to the expected mass of a dimer (232 kDa) than that of a monomer.

The apparent overestimation of TnpA MW could reflect some degree of flexibility or extended conformation in solution. To check this, we performed a dimensionless Kratky (dKratky) plot analysis to compare SAXS data obtained for TnpA^WT^ with the data obtained for a globular (BSA) or highly flexible (hTau40wt) protein (*SI Appendix*, Fig. S1*D*). The results suggest that TnpA^WT^ exists as a rather compact molecule in solution.

High-resolution cryo-EM structures of apo TnpA^WT^ and DNA-bound TnpA^S911R^ mutant were determined (31) which showed that TnpA is a stable dimer with 2-fold symmetry in both states. To further support relevance of these conclusions for the mutants, cryo-EM data were collected from TnpA^S911R^ in apo form and from TnpA^3X^ in complex with 48 bps long transposon ends (Fig. 1*B*, Fig. S2 and Table S2). We recorded a small dataset for apo TnpA^S911R^, which was enough to obtain 2D class averages showing shape and size similarity with apo TnpA^WT^ suggesting that TnpA^S911R^ is a dimer in apo state. Moreover, 6.1 Å 3D reconstruction calculated for TnpA^3X^-IR48 complex revealed a dimeric structure similar to TnpA^S911R^-IR48 indicating that the mutations do not change organization or conformation of the complexes (Fig. 1*B*).

Together with the SEC-SAXS analysis, the cryo-EM data support the conclusion that free TnpA wild type and hyperactive mutants exists as a ‘closed’ dimer in solution, and that PEC formation involves a substantial conformational change that allows TnpA to bind tightly two transposon ends in an active configuration. No evidence of single-occupancy SEC complex could be found in the TnpA^S911R^-IR48 and TnpA^3X^-IR48 cryo-EM datasets, which is consistent with DNA binding by the two mutant TnpAs being highly cooperative, showing much higher affinity for two transposon ends than for one [(17), see also below]. Only a few particles corresponding to singly bound proteins were observed when TnpA^WT^ was imaged in the presence of an IR-containing DNA fragment (31). However, compared to free protein, the fraction of particles corresponding to putative SEC conformation was too low for calculating 3D reconstructions from them. This likely reflects the low affinity of SEC relative to PEC, as revealed by EMSA experiments (17).

### TnpA binding to a single transposon end

According to previous biochemical studies (17), SEC formation appears to be a transient stage in the assembly of the transposition complex, with the subsequent structural transition leading to PEC being a rate-limiting step in the transposition process (Fig. 1*A*). AFM force spectroscopy has been used to evaluate the formation and stability of SEC by comparing the binding strength of TnpA^WT^, TnpA^3X,^ and TnpA^S911R^ to DNA substrates containing a single Tn4430 end. Using ‘click’ chemistry (42), short linear DNA molecules carrying an alkyne reactive group at their 5’ end (*SI Appendix*, Table S4) were covalently immobilized in an oriented manner onto gold surfaces coated with alkanethiols terminated with polyethyleneglycol (PEG) with a small fraction of them (5%) harboring an azide group (PEG-N_3_) (Fig. 2*A*). These model surfaces were imaged by AFM, and the thickness of the grafted layer was validated by a scratching experiment, revealing a deposited layer of ∼1.5 and 2 nm for PEG/PEG-N_3_ and PEG/PEG-N_3_/one end respectively (*SI Appendix*, Fig. *S3A-E*).

**Fig 2.**
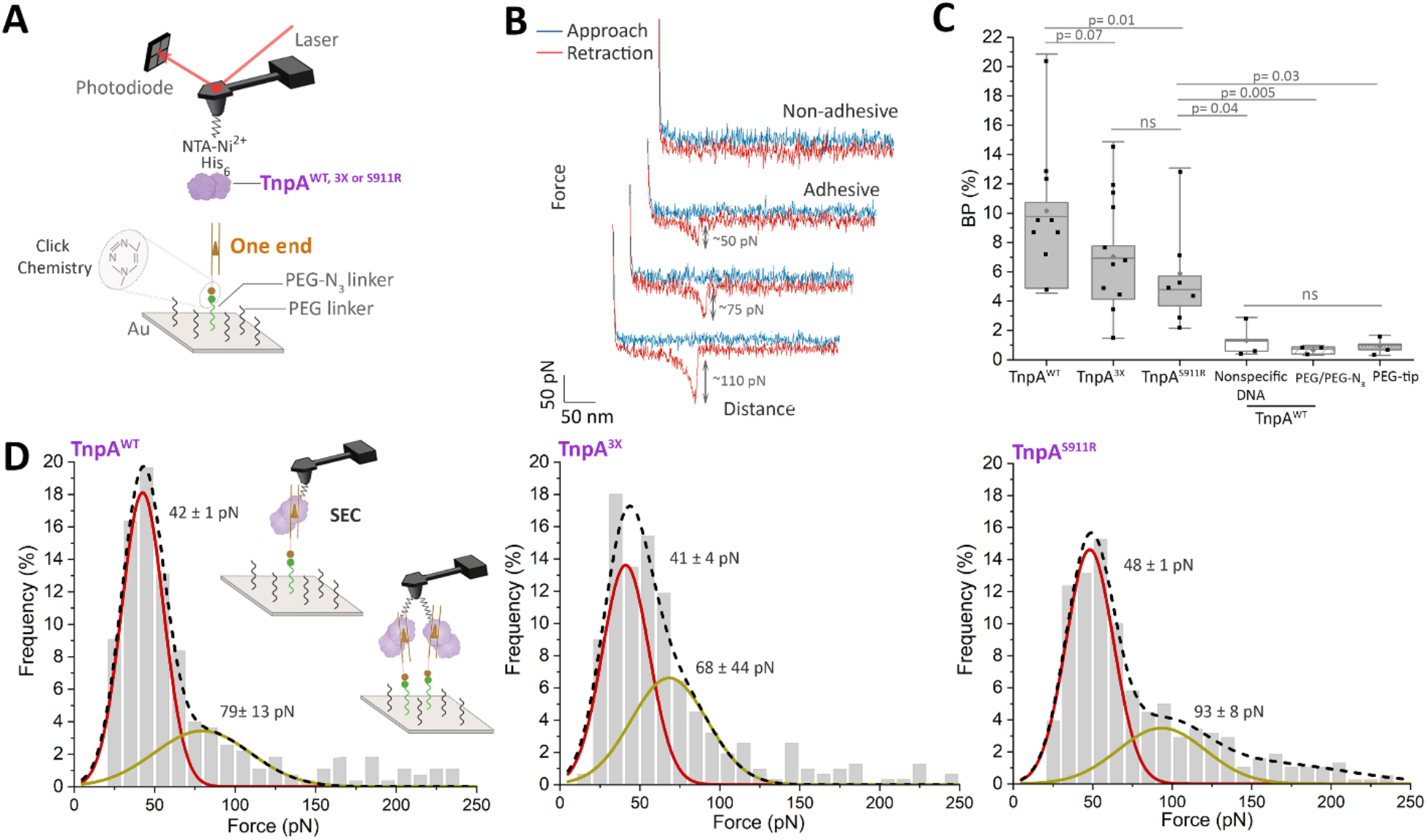
Probing of TnpA binding to linear DNA containing a single transposon end. (A) AFM set up. The interactions between TnpA and DNA are monitored by AFM on model surfaces. DNA molecules containing one transposon ends are attached to a surface using click chemistry (34). The AFM tip is functionalized by PEG linkers to which Ni^2+^-loaded NTA (Ni^2+^-NTA) is bound. TnpA contains a His6-tag (purple triangle), which can specifically bind the Ni^2+^-NTA ligands. (B) Representative force-distance curves showing either non-adhesive or specific adhesive interactions. The rupture force measured in each case is indicated to the right of the curves. (C) Box plot of AFM-measured binding probability (BP) between tips functionalized with TnpA^WT^, TnpA^3X,^ or TnpA^S911R^ as indicated, and surfaces grafted with DNA molecules containing one transposon end. The binding frequencies between TnpA^WT^ and the nonspecific DNA molecule or solely the PEG/PEG-N3 layer are shown as a specificity control. The binding probability measured with a TnpA-free polyethylene glycol (PEG) tip and the one-end surface gives the background level of the experimental setup. The horizontal line within the box indicates the median, the boundaries of the box indicate the 25th and 75th percentile, and the whiskers indicate the highest and lowest values of the results. The gray square in the box indicates the mean. For all experiments, data are representative of at least N=3 independent experiments. P values were determined by a two-sample t-test with R software (version 3.1.1). (D) Frequency distributions of rupture forces between TnpA^WT^, TnpA^3X,^ and TnpA^S911R^ and the one-end substrate. The red curve corresponds to SEC while the brown curve probably corresponds two interactions established in parallel. The dashed line corresponds to the sum of the red and brown curves.

To study the binding of TnpA to DNA, purified TnpA^WT^, TnpA^3X,^ and TnpA^S911R^ proteins were attached in an oriented manner to tips AFM functionalized with Ni^2+^-NTA groups through the His-tag present at the C-terminus (33). Force-distance (FD) curves were recorded to probe the binding strength of the three TnpA variants to the one-end DNA substrate (Fig. 2*A,B*). Specific adhesion events were observed in 10.7 ± 4.5% (mean ± S.D., *N*= 18845), 7.7 ± 4.1% (*N*= 22548) and 5.7 ± 3.7% (*N*= 14336) of the retraction FD curves for TnpA^WT^, TnpA^3X^ and TnpA^S911R^, respectively (Fig. 2*C*).

To confirm the specificity of these interactions, independent control experiments were performed using (*i*) a surface functionalized with nonspecific DNA (sequence shown in *SI Appendix*, Fig. S4), (*ii*) a surface-functionalized only with the PEG/PEG-N_3_ mix (without DNA), and (*iii*) an AFM tip functionalized only with the PEG thiol. As expected, TnpA did not bind significantly to non-specific DNA nor PEG, and the binding frequency was also significantly reduced when the AFM tip was functionalized with the PEG linker only (Fig. 2*C*). Finally, a competition assay was performed by injecting increasing amounts of a free DNA molecule carrying the Tn4430IR end sequence within the AFM reaction chamber (*SI Appendix*, Fig. S5). For all three TnpA proteins, a reduction of the binding probability (BP) between the immobilized partners was observed as a function of the concentration of the competitor DNA. Altogether, these controls confirm the specificity of the interactions and the critical importance of the IR sequence in the formation of SEC.

Next, we extracted the binding strength from the individual FD curves and analyzed the force distribution of the interactions established between the one-end substrate and the three TnpA proteins. The binding forces were displayed as a histogram for each TnpA variant and we observed a peak in the range 40-50 pN in each case (Fig. 2*D*). We attributed this interaction to the formation of the SEC. In addition, at a lower frequency, the second population of interactions is present in the range 70-95 pN, which likely indicates the formation of two interactions established in parallel (Fig. 2*D*), presumably between two different TnpA molecules attached to the AFM tip and two DNA fragments present on the surface.

These results show that TnpA^WT^, TnpA^3X,^ and TnpA^S911R^, can bind to a single transposon end to form the SEC and that at a first sight, the primary determinants of this interaction are the same for the three proteins as was demonstrated by DNA footprinting analysis (17).

### Exploring the kinetics properties of SEC

To characterize the binding properties of the interactions occurring between TnpA and the one-end substrate, the binding strength of the complex was probed at various force loading rates (LR; *i*.*e*. by increasing the force acting on the bond over time), enabling to extract of its kinetic parameters (43). We approximate the binding free-energy landscape as a simple two-state model in which the bound state is separated from the unbound state by a single energy barrier (Fig. 3*A*). Dynamic force spectroscopy (DFS) plots were obtained for each TnpA-one end system (Fig. 3*B-D*). In each case, the unbinding force increased linearly with the logarithm of the LR. Briefly, the individual rupture forces (every single gray data point in Fig. 3*B-D*) were extracted along with their loading rate, which is estimated from the slope of the force *vs* time curve just before the rupture, as previously established (44). Then, bond strengths were analyzed through distinct discrete ranges of LRs, plotted as force histograms, and further fitted with a multipeak Gaussian distribution (45). This analysis allowed us to determine for each discrete range of LR the most probable unbinding force for the single or multiple interactions (Fig. 3*B-D*, black squares). Afterward, the Bell-Evans (BE) model (37, 38) was used to fit the single-rupture event, providing access to the kinetics parameters (Fig. 3*B-D*, solid line). First, the width of the energy barrier (*x*_u_) was extracted from the slope of the fit. We obtained the following *x*_u_ values: 0.74 ± 0.05 nm, 0.79 ± 0.09 nm and 0.67 ± 0.11 nm for TnpA^WT^, TnpA^3X^ and TnpA^S911R^, respectively. These values confirm that the bonds are similar for the three systems and are characteristic of a stiff bonding network (46). The kinetic off-rate (*k*_off_) or dissociation rate obtained from the intersection of the fit (at LR =0) yielded *k*_off_ values of 0.20 ± 0.06 s^−1^, 0.10 ± 0.10 s^−1^, and 0.10 ± 0.14 s^−1^ for TnpA^WT^, TnpA^3X^, and TnpA^S911R^, respectively (Fig. 3*B-D*). These values are within the range of those reported for the LexA repressor (which controls approximately 31 genes involved in the SOS response of *Escherichia coli*) as determined by AFM-based single-molecule force spectroscopy (46). The LexA repressor showed dissociation rate constants of 0.045 s^-1^ and 0.13 s^-1^ for the *recA* and *yebG* operons, respectively (46).

**Fig 3.**
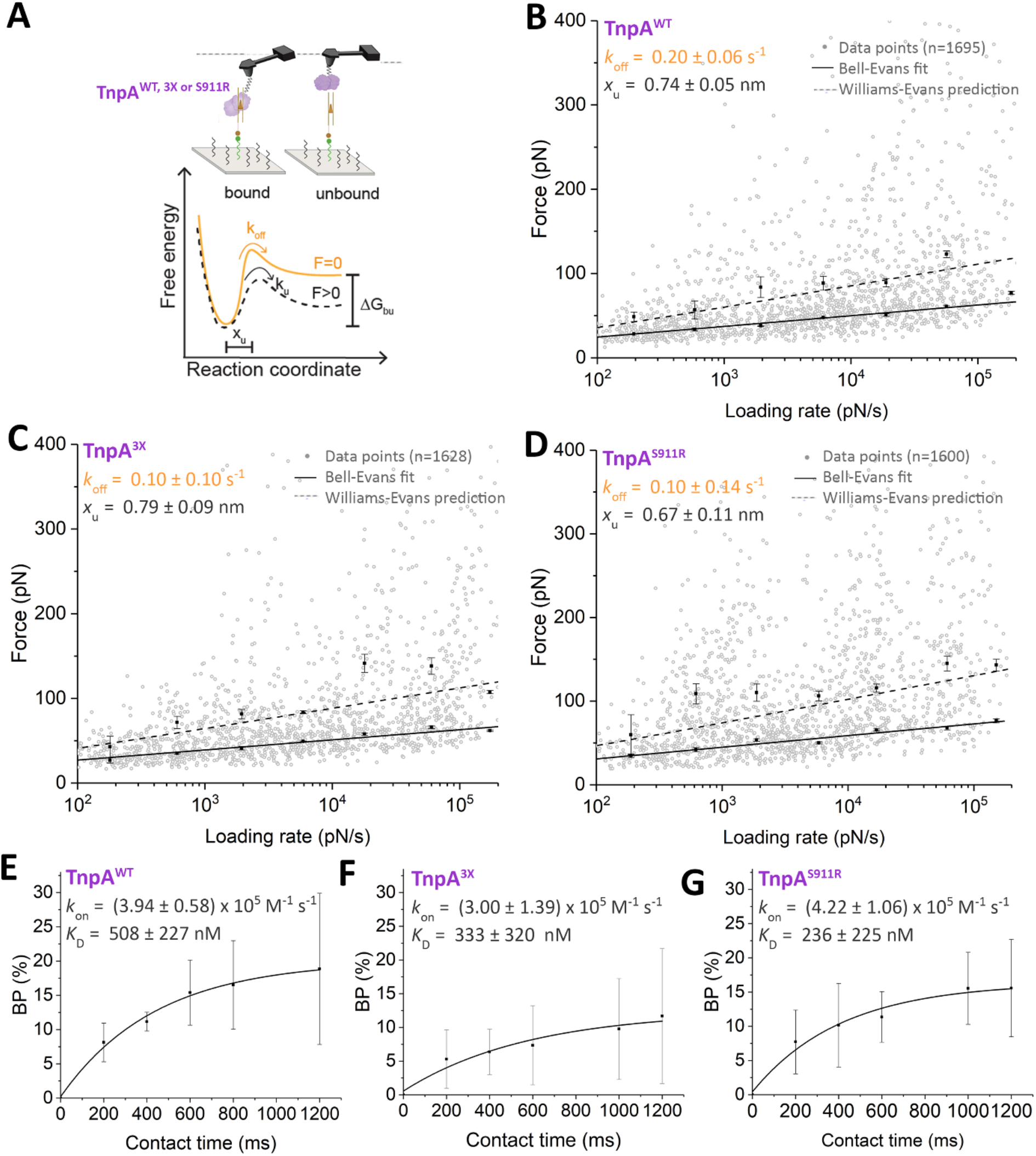
Dynamics of SEC. (A) The binding of TnpA^WT^, TnpA^3X,^ or TnpA^S911R^ is probed on a surface coated with one-end DNA molecules. Bell–Evans model describes a ligand-receptor bond as a simple two-state model. The bound state is separated from the unbound state by an energy barrier located at distance *X*u. *K*u and *K*_off_ represent the transition rate and transition at thermal equilibrium, respectively. (B-D) Dynamic force spectroscopy (DFS) plot showing the distribution of the rupture forces as a function of the loading rates measured either between TnpA^WT^ (B), TnpA^3X^ (C) or TnpA^S911R^ (D) and the one-end-coated surface. Data corresponding to single interactions are fitted with the Bell-Evans (BE) model (I, black curve). Dashed lines represent the predicted binding forces for two simultaneous uncorrelated interactions (Williams-Evans model). Error bars indicate standard deviations (SD) of the mean values. Experiments were reproduced at least five times with independent tips and samples. (E-G) The binding probability (BP) is plotted as a function of the contact time for TnpA^WT^ (E), TnpA^3X^ (F) or TnpA^S911R^ (G) on the one end-coated surface, and data points were fitted using a least-squares fit of a monoexponential growth. One data point belongs to the BP from one map acquired at 1 μm/s retraction speed for the different contact times. Experiments were reproduced at least three times with independent tips and samples. Error bars indicate s.d. of the mean values.

Using the predictive Williams-Evans (WE) model (39) we analyzed the higher force observed on the DFS plot. The second peaks observed at each loading rate can be attributed to uncorrelated bonds loaded in parallel (Fig. 3*B-D*, dashed line), confirming that under our experimental conditions two SEC complexes can be simultaneously probed.

### Approaching the binding-free energy landscape

Assuming that SEC can be approximated by pseudo-first-order kinetics, the kinetic on-rate (*k*_on_) was estimated from our single-molecule force spectroscopy experiments by monitoring the binding probability (BP) between the different pairs of partners at various contact times (Fig. 3*E-G*). k_on_ values were extracted by taking the effective concentration as the number of binding partners (TnpA-one end) within an effective volume *V*_eff_ accessible under free-equilibrium interaction. *V*_eff_ can be approximated by a sphere whose radius includes the linker, the TnpA protein, and the linear one-end DNA substrate (see equations (1) and (2) in materials and methods). The binding frequency follows a logarithmic growth with the contact time for all three TnpAs. The values extracted for interaction time were 0.44 ± 0.06 s, 0.58 ± 0.27 s and 0.41 ± 0.10 s, which led to a *k*_on_ of 3.94 × 10^5^ M^−1^ s^−1^, 3.00 × 10^5^ M^−1^ s^−1^ and 4.22 × 10^5^ M^−1^ s^−1^ for TnpA^WT^, TnpA^3X,^ and TnpA^S911R^ complexes, respectively. Finally, the dissociation constant *K*_D_, calculated as the ratio between *k*_off_ and *k*_on_ yielded values around ∼508 nM, 333 nM, and 236 nM for the TnpA^WT^, TnpA^3X,^ and TnpA^S911R^ complexes, respectively (Fig. 3*E-G*). For single-molecule interactions, the bond lifetime τ can be directly related to the inverse kinetic off-rate (τ = *k*_off_^−1^), resulting here in a τ of 5 s for TnpA^WT^ and 10 s for both hyperactive mutants. These values correspond to high-affinity interactions (47), which confirms the specificity of the complexes established between the different TnpA proteins and the one-end substrate.

### TnpA binding to two transposon ends and PEC formation

According to our model, PEC formation is a critical step of the Tn4430 transposition pathway during which the TnpA dimer bound to a single transposon end undergoes a conformational change that allows it to capture the opposite end of the transposon and to adopt an active configuration (Fig. 1*A*). We believe that this is a rate-limiting step in the activation pathway, since TnpA^WT^ hardly forms the PEC under standard conditions, whereas for the hyperactive TnpA^3X^ and TnpA^S911R^ mutants, PEC formation is highly cooperative (17). However, so far, the dynamics of PEC formation is poorly understood due to the lack of direct quantitative information.

To this end, a specific DNA substrate containing inversely oriented Tn4430 IR ends was assembled by annealing three complementary oligonucleotides (*SI Appendix*, Fig. S6*A*, and Table S4). The duplex regions corresponding to the IR ends are separated by a 20-nt single-stranded DNA linker to allow for enough flexibility to pair both transposon ends inside the PEC. Restriction digest followed by gel electrophoresis showed that ∼100% of the DNA substrates assembled correctly (*SI Appendix*, Fig. S6*B*). The central position of the DNA linker was substituted with an alkyne group-containing dU nucleotide to anchor the substrate to azide-modified AFM surfaces by click chemistry (42) (Fig. 2*A* and Fig. 4*B*). This model surface was imaged by AFM, and the thickness of the grafted layer was validated by a scratching experiment, revealing a deposited layer of 2 nm (*SI Appendix*, Fig. S6*C,D*). On the other hand, purified TnpA proteins were attached to the AFM tip as described above for the one-end experiments.

**Fig 4.**
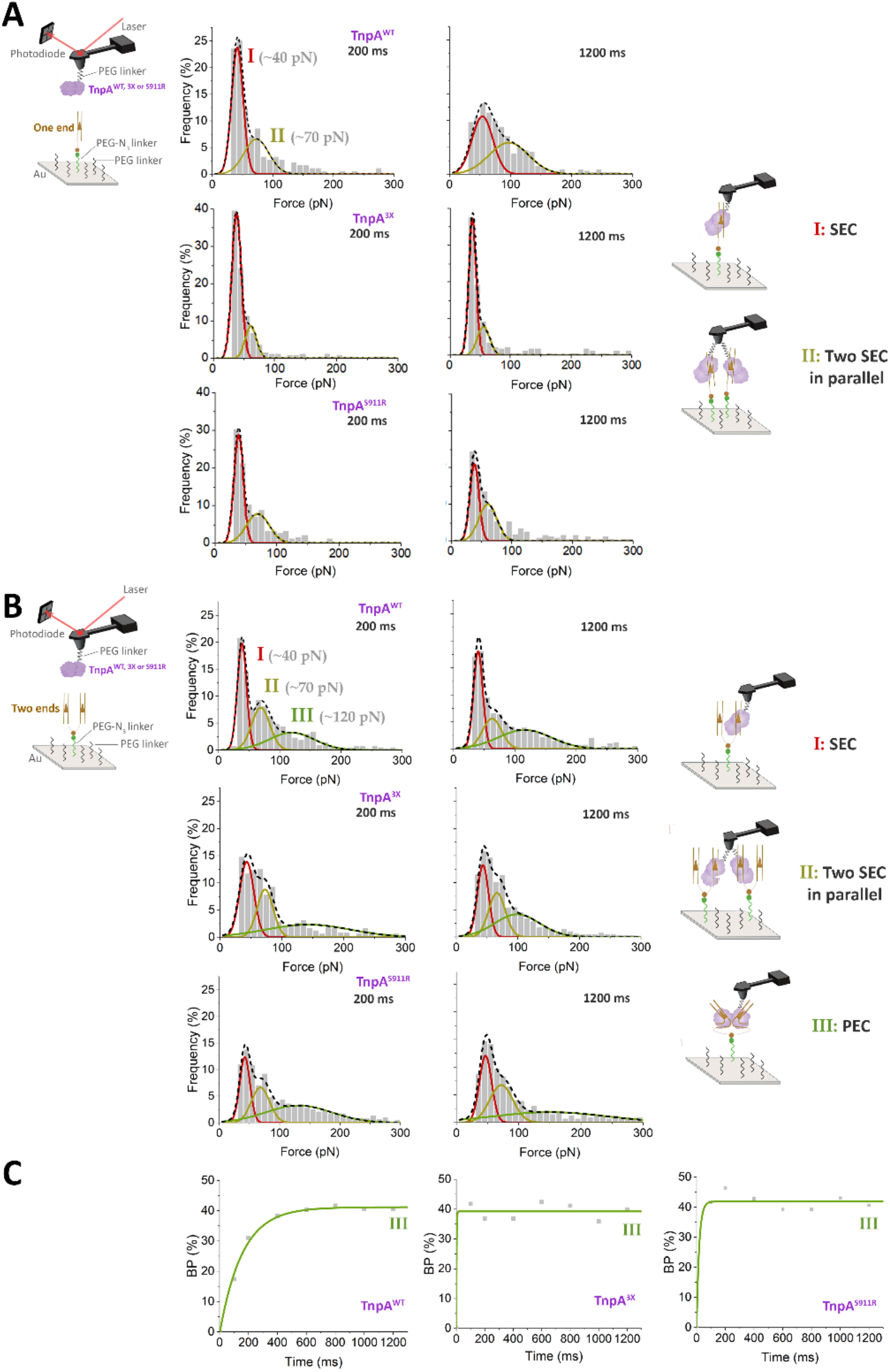
Probing PEC formation. (A and B) Binding of TnpA to surface coated with one-end molecules is compared to TnpA binding to two transposon ends. Frequency distributions of rupture forces at 200 ms and 1200 ms of contact time between either TnpA^WT^, TnpA^3X^ or TnpA^S911R^, and the one-end substrate (A) or the two-end substrate (B) were extracted from data presented in *SI Appendix*, Fig. S7, S8 and S9. I, II or III populations are shown by red, brown or green curves respectively, and their corresponding interpretation is schematized to the right of each panel. (C) Binding probability (BP) as a function of contact time for the third population (III).

Conformational dynamics occurring during PEC formation were analyzed by FD-based AFM by tracking the binding strength between the various transposases and the substrate with either one end (Fig. 4*A*) or two ends (Fig. 4*B*). To follow the dynamics, the contact time between the binding partners was progressively increased from 200 to 1200 ms, to provide sufficient time for binding and conformational transition (*SI Appendix*, Fig. S7-S9).

While only two populations are observed with the one-end substrate (Fig. 4*A*), a third population at higher forces appeared in the histograms for the two-end substrate (Fig. 4*B*) and the three TnpA. As evidenced above, most probed interactions were monomolecular (*i*.*e*., between one TnpA molecule and a single IR end) with an average force of ∼40 pN corresponding to SEC formation (Fig. 4*A,B*, *scheme I*). A second sub-population with a force of ∼70 pN was observed confirming the possibility of forming two SEC in parallel with both the one-end and two-end substrates (Fig. 4*A,B*, *scheme II*). Interestingly, only in the experiments with the two-end substrate, a third population around ∼120 pN was observed. Since this third population only appears with the two-end substrate, we attributed it to the formation of a complex between a single TnpA molecule bound to both transposon ends of the substrate, thus mimicking PEC formation (Fig. 4*B*, *scheme III*). For the three TnpAs, the strength of interactions measured for the third peak is higher than that measured for parallel interactions between TnpA and two independent ends, supporting that PEC formation involves a larger network of contacts between TnpA and DNA than SEC. Looking at the dynamics (Fig. 4*C*), we observed that PEC formation by TnpA^WT^ follows pseudo-first-order association kinetics with a time constant (τ) of ∼156 ms. By contrast, for the TnpA^3X^ and Tnp^S911R^ hyperactive mutants, the kinetic analysis showed that complex III formation occurred at a much faster rate (τ of 2 ms and 17 ms, respectively), at the limit of time resolution of the experiment.

## DISCUSSION

Transposition, like other complex protein-DNA transactions, is a carefully orchestrated process involving multiple partners. In particular, transposition catalyzed by DDE/D transposases proceeds through a series of DNA cleavages and strand transfer reactions fusing the transposon ends to a new target locus (2). Since these reactions are potentially harmful to the host, it is important that each step be irreversible and coordinated with each other. Directionality is generally thought to result from energetically favorable conformational changes taking place in the transposition complex to rearrange and adequately position the partners at each step of the reaction (14, 48–51). In many cases, initial activation of the reaction results from the formation of an oligomeric nucleoprotein complex termed ‘transposome’ in which both transposon ends are brought together by the transposase. In the simplest situation referred to as ‘synapsis-by-protein-dimerization’ (S-PD), the transposase separately binds as a monomer to each transposon end and then oligomerizes to assemble the synaptic complex (52). However, together with previous biochemical data, the finding that the TnpA transposase of Tn4430 forms a stable dimer in solution indicates that it uses a conceptually different ‘synapsis-by-naked-end-capture’ (S-NEC) mechanism in which an inactive TnpA dimer first bind to a single transposon end to form the SEC and then undergoes a conformational change to capture the other end and adopt the active PEC configuration (52) (Fig. 1 and Fig. 5).

**Fig 5.**
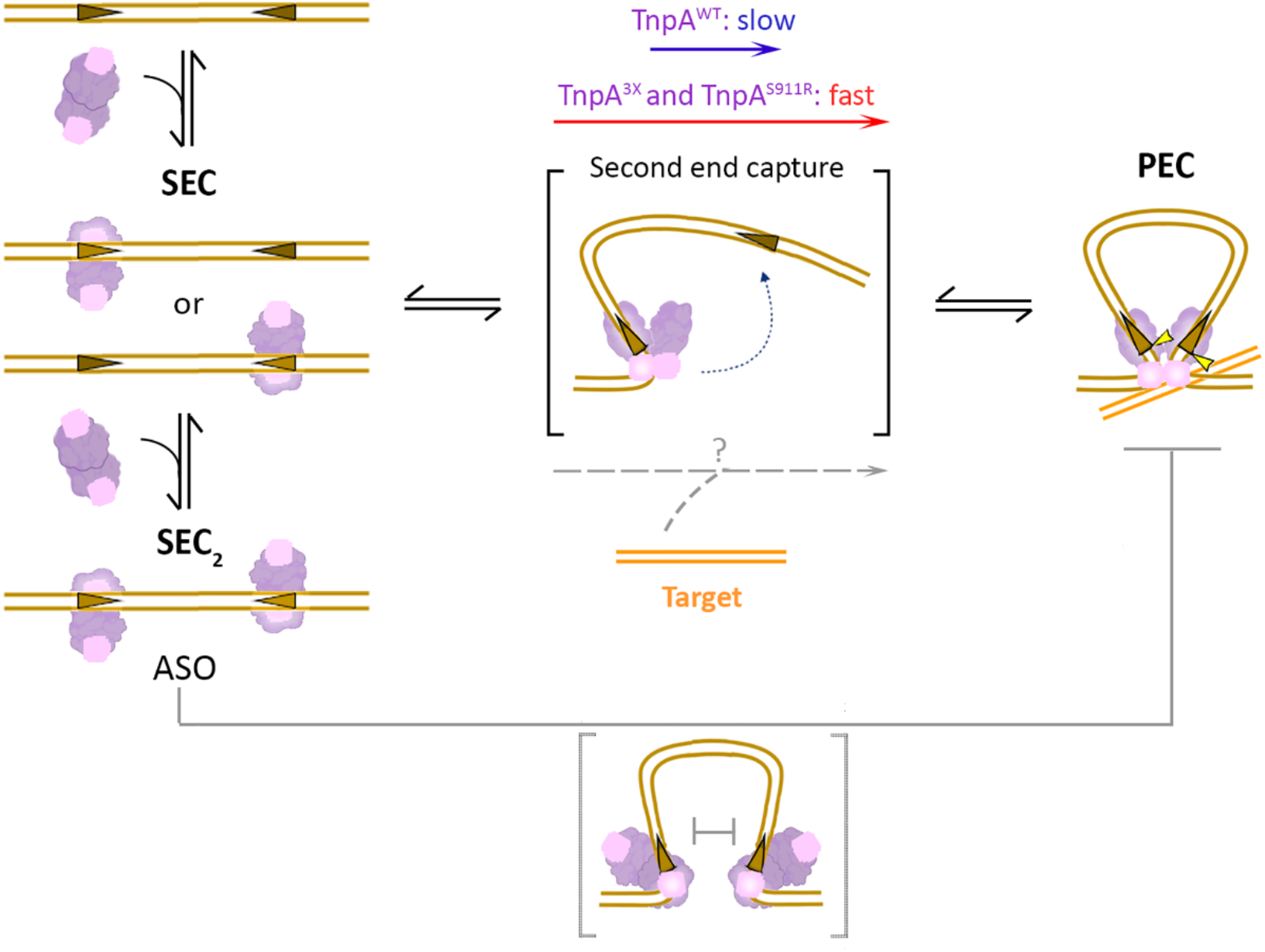
Model for Tn4430 transpososome activation pathway. Synapsis-by-naked-end-capture (S-NEC) mechanism in which an inactive TnpA dimer first bind to single transposon end to form the inactive SEC and then undergoes a conformational change to capture the other end and adopt the active PEC configuration. ASO: assembly site occlusion. See text for further details.

Here, we have used FD-based AFM to decipher the dynamic of Tn4430 transposition complex formation at the single particle level by comparing the activities of TnpA^WT^ with that of the two hyperactive variants TnpA^3X^ and TnpA^S911R^, which are thought to reproduce the two key intermediates of the assembly pathway. We took advantage of click chemistry (42) to assemble and decorate the AFM surfaces with specifically designed one-end and two-end DNA substrates allowing us to collect high throughput data for both the SEC and PEC with high selectively and reproductivity, an approach that could readily apply to any other protein-DNA system. The results show that TnpA^WT^ and both hyperactive mutants specifically bind to a spatially constrained (i.e., non-diffusible) single transposon end with quite similar kinetic and thermodynamic parameters, indicating that the determinants for SEC formation are essentially the same for the three proteins. The measured force of interaction (∼45 pN) and the deduced *k*_on_ (∼ 3 to 4 10^5^ M^-1^ s^-1^) are typical of specific high-affinity bimolecular associations such as between a protein-ligand and its receptor or an enzyme with its substrate, which is consistent with [(OP)2-Cu+] footprint patterns showing that within the SEC, both the wild type and the mutant TnpAs protect the same ∼30-bp region at the inner part of Tn4430 38-bp IR sequence, making specific contacts on both DNA strands (17).

However, the SEC appears to be a metastable intermediate of transpososome assembly, with an estimated lifetime τ being comprised between 5 s for TnpA^WT^ and 10 s for TnpA^3X^ and TnpA^S911R^, respectively. As discussed below, reversible binding of TnpA to the transposon ends is important since it gives the possibility to regulate the initial step of transposition prior to proceeding further by forming the PEC or not (Fig. 5). The two-fold lower *k*_off_ measured for TnpA^3X^ and TnpA^S911R^ compared to TnpA^WT^ (*i*.*e*., 0.1 s^-1^ *vs* 0.2 s^-1^) suggests that both hyperactive mutants have a higher propensity to adopt a transitional state for PEC formation, even in the absence of the partner end.

Supporting this view, kinetic analysis performed with a two-end substrate showed that PEC formation by TnpA^3X^ and TnpA^S911R^ occurs at a much faster rate than with TnpA^WT^, being practically instantaneous. This result is consistent with previous EMSA titration experiments showing that PEC formation by TnpA^3X^ and TnpA^S911R^ is highly cooperative, with the SEC being not or barely detectable even at low protein-DNA ratios (17) and with cryo-EM datasets showing that incubation of both deregulated TnpA mutants with IR DNA substrates almost exclusively gives PEC (Fig. 1*B*) (31). The higher force required to disrupt the PEC compared to two independent SEC (*i*.*e*., ∼120 pN *vs* ∼70-80 pN) is also consistent with DNA footprint analyses and high-resolution structural data showing that TnpA binding into the PEC makes extended contacts with the outer part of the transposon end to activate the transposon cleavage sites (17, 21).

Thus, together the results support the conclusion that SEC to PEC transition is a rate-limiting step in the Tn4430 S-NEC assembly pathway, and that specific mutations in TnpA^3X^ and TnpA^S911R^ have lowered the energy barrier for PEC formation, making the hyperactive mutants more prone than TnpA^WT^ to undergo the required conformational change to capture both tranposon ends and become catalytically active (Fig. 5) (31).

What then promotes activation of TnpA under the wild-type situation? Since TnpA^3X^ and TnpA^S911R^ were initially isolated for their impairment in transposition immunity, they are less demanding than TnpA^WT^ concerning target selection, being able to transpose at a higher frequency into non-permissive DNA molecules (13). We, therefore, propose that SEC to PEC transition is normally controlled by the target, with permissive targets providing a positive signal to induce the formation of PEC and/or immune targets preventing the activating conformational change (Fig. 5). This target activation checkpoint would ensure that all partners are brought together before triggering transposition and provide a rationale for target immunity and the behavior of immunity-defective TnpA mutants. The presence of a proper target may either promote SEC to PEC transition or stabilize the PEC in the active configuration (Fig. 5). We favor the latter possibility since TnpA^WT^ was shown to form the PEC on the geometrically favorable two-end substrate, albeit with lower efficiency than TnpA^3X^ and TnpA^S911R^.

The metastability of SEC and the implication of a putative activation mechanism to stabilize the PEC should also provide a means to counteract a process known as ‘assembly site occlusion’ (ASO) that is proposed to act as an autoregulation mechanism to limit the activity of transposons, mostly from eukaryotes, which like Tn4430 transpose via the S-NEC pathway (52). In this mechanism, termed ‘overproduction inhibition’ (OPI), increasing expression of the transposase dimer resulting from the genomic expansion of the element gradually saturate the transposon ends, thereby creating a barrier for PEC assembly. There is currently no evidence for an OPI regulatory mechanism acting on Tn4430 transposition in vivo. In vitro, the PEC assembled by the hyperactive TnpA^3X^ and TnpA^S911R^ mutants was only shown to dissociate into SECs when a DNA substrate was incubated with over-saturating concentrations of the protein (17). We propose that under favorable conditions for active transpososome assembly (*e*.*g*., in presence of an appropriate target), stabilization of the PEC would displace the doubly bound SEC_2_ state, thereby pushing the reaction forward (Fig. 5). To validate this model, our current work aims at characterizing which properties of the target are involved in integration site discrimination by Tn4430. Determination of the structure of TnpA and the TnpA-DNA complexes will also highlight the molecular determinants of the SEC to PEC transition and the role of the target in the activation mechanism.

## DATA AVAILABILITY

All study data are included in the article and/or supporting information. The AFM data have been deposited in Figshare (https://figshare.com/articles/figure/Fernandez_Source_Data_xlsx/20032232). The SAXS data were deposited in the small-angle scattering biological data bank under accession code SASDMR5. The cryo-EM maps of the TnpA 3x-IR48 and TnpA WT have been deposited with accession codes EMD-15213 and EMD-15218.

## Author Contributions

M.F., B.H., and D.A. conceived the project, planned the experiments, and analyzed the data. M.F conducted the AFM experiments. C.S. and S.D. provided technical support in protein purification and AFM manipulation, respectively. A.V.S., R.G.E. and Y.L. conducted and analyzed the SAXS and Cryo-EM experiments. M.F., B.H. and, D.A. wrote the draft manuscript. All authors read and approved the manuscript.

We thank Gopinath <u>Muruganandam</u> for his help with SAXS data collection. The authors are grateful for beam time and excellent beamline support at SWING beamline of SOLEIL Synchrotron.

## REFERENCES

1. Craig, N. (2015) A Moveable Feast: An Introduction to Mobile DNA. In Mobile DNA III.pp. 3–39.

2. Hickman, A.B. and Dyda, F. (2016) DNA Transposition at Work. Chem. Rev., 116, 12758–12784.

3. Aziz, R.K., Breitbart, M. and Edwards, R.A. (2010) Transposases are the most abundant, most ubiquitous genes in nature. Nucleic Acids Res., 38, 4207–4217.

4. Kazazian, H.H. (2004) Mobile Elements: Drivers of Genome Evolution. Science., 303, 1626–1632.

5. Frost, L.S., Leplae, R., Summers, A.O. and Toussaint, A. (2005) Mobile genetic elements: The agents of open source evolution. Nat. Rev. Microbiol., 3, 722–732.

6. Wells, J.N. and Feschotte, C. (2020) A Field Guide to Eukaryotic Transposable Elements. Annu Rev Genet., 54, 539–561.

7. Guy-Franck, R. (2020) Eukaryotic Pangenomes. In Tettelin, H., Medini, D., Editors (eds), The Pangenome: Diversity, Dynamics and Evolution of Genomes. Cham (CH): Springer.

8. Partridge, S.R., Kwong, S.M., Firth, N. and Jensen, S.O. (2018) Mobile genetic elements associated with antimicrobial resistance. Clin. Microbiol. Rev., 31, 1–61.

9. Nicolas, E., Lambin, M., Dandoy, D., Galloy, C., Nguyen, N., Oger, C.A. and Hallet, B. (2015) The Tn3 family of Replicative Transposons. Microbiol. Spectr., 3, 1–32.

10. Cuzon, G., Naas, T. and Nordmann, P. (2011) Functional characterization of Tn4401, a Tn3-based transposon involved in bla KPC gene mobilization. Antimicrob. Agents Chemother., 55, 5370–5373.

11. Snesrud, E., Maybank, R., Kwak, Y.I., Jones, A.R., Hinkle, M.K. and McGann, P. (2018) Chromosomally Encoded mcr-5 in Colistin-Nonsusceptible Pseudomonas aeruginosa. Antimicrob. Agents Chemother., 62, e00679–18.

12. Brandt, C., Viehweger, A., Singh, A., Pletz, M.W., Wibberg, D., Kalinowski, J., Lerch, S., Müller, B. and Makarewicz, O. (2019) Assessing genetic diversity and similarity of 435 KPC-carrying plasmids. Sci. Rep., 9, 11223.

13. Lambin, M., Nicolas, E., Oger, C.A., Nguyen, N., Prozzi, D. and Hallet, B. (2012) Separate structural and functional domains of Tn4430 transposase contribute to target immunity. Mol. Microbiol., 83, 805–820.

14. Montaño, S.P. and Rice, P.A. (2011) Moving DNA around: DNA transposition and retroviral integration. Curr Opin Struct Biol., 21, 370–378.

15. Nowotny, M. (2009) Retroviral integrase superfamily: the structural perspective. EMBO Rep., 10, 144–151.

16. Majorek, K.A., Dunin-Horkawicz, S., Steczkiewicz, K., Muszewska, A., Nowotny, M., Ginalski, K. and Bujnicki, J.M. (2014) The RNase H-like superfamily: New members, comparative structural analysis and evolutionary classification. Nucleic Acids Res., 42, 4160–4179.

17. Nicolas, E., Oger, C.A., Nguyen, N., Lambin, M., Draime, A., Leterme, S.C., Chandler, M. and Hallet, B.F.J. (2017) Unlocking Tn3-family transposase activity in vitro unveils an asymetric pathway for transposome assembly. Proc. Natl. Acad. Sci. U. S. A., 114, 669–678.

18. Guynet, C., Nicolas, E., Ton-Hoang, B., Bouet, J. and Hallet, B. (2020) First biochemical steps on bacterial transposition pathways. Methods Mol Biol., 2075, 157–177.

19. Nicolas, E., Lambin, M. and Hallet, B. (2010) Target immunity of the Tn3-family transposon Tn4430 requires specific interactions between the transposase and the terminal inverted repeats of the transposon. J. Bacteriol., 192, 4233–4238.

20. Müller, D.J., Dumitru, A.C., Lo Giudice, C., Gaub, H.E., Hinterdorfer, P., Hummer, G., De Yoreo, J.J., Dufrêne, Y.F. and Alsteens, D. (2020) Atomic Force Microscopy-Based Force Spectroscopy and Multiparametric Imaging of Biomolecular and Cellular Systems. Chem. Rev.

21. David, G. and Pérez, J. (2009) Combined sampler robot and high-performance liquid chromatography: a fully automated system for biological small-angle X-ray scattering experiments at the Synchrotron SOLEIL SWING beamline. J. Appl. Crystallogr., 42, 892–900.

22. Shkumatov, A. V. and Strelkov, S. V. (2015) DATASW, a tool for HPLC-SAXS data analysis. Acta Crystallogr. D. Biol. Crystallogr., 71, 1347–1350.

23. Durand, D., Vivès, C., Cannella, D., Pérez, J., Pebay-Peyroula, E., Vachette, P. and Fieschi, F. (2010) NADPH oxidase activator p67phox behaves in solution as a multidomain protein with semi-flexible linkers. J. Struct. Biol., 169, 45–53.

24. Svergun, D.I. (1992) Determination of the regularization parameter in indirect-transform methods using perceptual criteria. J. Appl. Crystallogr., 25, 495–503.

25. Svergun, D. (1999) Restoring low resolution structure of biological macromolecules from solution scattering using simulated annealing. Biophys. J., 76, 2879–2886.

26. Efremov, R.G. and Stroobants, A. (2021) Coma-corrected rapid single-particle cryo-EM data collection on the CRYO ARM 300. Acta Crystallogr. Sect. D Struct. Biol., 77, 555–564.

27. Mastronarde, D.N. (2005) Automated electron microscope tomography using robust prediction of specimen movements. J. Struct. Biol., 152, 36–51.

28. Zivanov, J., Nakane, T., Forsberg, B.O., Kimanius, D., Hagen, W.J.H., Lindahl, E. and Scheres, S.H.W. (2018) New tools for automated high-resolution cryo-EM structure determination in RELION-3. Elife, 7, 1–22.

29. Zheng, S.Q., Palovcak, E., Armache, J.-P., Verba, K.A., Cheng, Y. and Agard, D.A. (2017) MotionCor2 - anisotropic correction of beam-induced motion for improved cryo-electron microscopy. Nat Methods., 14, 331–332.

30. Rohou, A. and Grigorieff, N. (2015) CTFFIND4: Fast and accurate defocus estimation from electron micrographs. J. Struct. Biol., 192, 216–221.

31. Shkumatov, A. V, Aryanpour, N., Oger, C.A., Goossens, G., Hallet, B.F. and Efremov, R.G. (2022) Metamorphism of catalytic domain controls transposition in Tn3 family transposases. bioRxiv.

32. Punjani, A., Rubinstein, J.L., Fleet, D.J. and Brubaker, M.A. (2017) CryoSPARC: Algorithms for rapid unsupervised cryo-EM structure determination. Nat. Methods, 14, 290–296.

33. Pfreundschuh, M., Alsteens, D., Wieneke, R., Zhang, C., Coughlin, S.R., Tampé, R., Kobilka, B.K. and Müller, D.J. (2015) Identifying and quantifying two ligand-binding sites while imaging native human membrane receptors by AFM. Nat. Commun., 6, 1–7.

34. Gruber, H.J. (2016) Functionalization of AFM tips with Click Chemistry.

35. Butt, H.-J. and Jaschke, M. (1995) Calculation of thermal noise in atomic force microscopy. Nanotechnology, 6, 1–7.

36. Bustamante, C., Marko, J.F., Siggia, E.D. and Smith, S. (1994) Entropic Elasticity of λ-Phage DNA. Science., 265, 1599–1600.

37. Evans, E. and Ritchie, K. (1997) Dynamic strength of molecular adhesion bonds. Biophys. J., 72, 1541–1555.

38. Evans, E.A. and Calderwood, D.A. (2007) Forces and bond dynamics in cell adhesion. Science., 316, 1148–1153.

39. Evans, E. & Williams, P. (2002) Dynamic force spectroscopy. In Physics of bio-molecules and cells. Physique des biomolécules et des cellules Ch. Springer, pp. 145–204.

40. Rankl, C., Wildling, L., Neundlinger, I., Kienberger, F., Bruber, H., Blaas, D. and Hinterdorfer, P. (2011) Determination of the Kinetic On- and Off-Rate of Single Virus–Cell Interactions. Methods Mol. Biol., 736, 197–210.

41. Rankl, C., Kienberger, F., Wildling, L., Wruss, J., Gruber, H.J., Blaas, D. and Hinterdorfer, P. (2008) Multiple receptors involved in human rhinovirus attachment to live cells. Proc. Natl. Acad. Sci. U. S. A., 105, 17778–17783.

42. Kolb, H.C., Finn, M.G. and Sharpless, K.B. (2001) Click Chemistry: Diverse Chemical Function from a Few Good Reactions. Angew. Chemie - Int. Ed., 40, 2004–2021.

43. Merkel, R., Nassoy, P., Leung, A., Ritchie, K. and Evans, E. (1999) Energy landscapes of receptor-ligand bonds explored with dynamic force spectroscopy. Nature, 397, 50–53.

44. Alsteens, D., Pfreundschuh, M., Zhang, C., Spoerri, P.M., Coughlin, S.R., Kobilka, B.K. and Muller, D.J. (2015) Imaging G protein–coupled receptors while quantifying their ligand-binding free-energy landscape. Nat. Methods, 12, 845–851.

45. Alsteens, D., Newton, R., Schubert, R., Martinez-Martin, D., Delguste, M., Roska, B. and Müller, D.J. (2017) Nanomechanical mapping of first binding steps of a virus to animal cells. Nat. Nanotechnol., 12, 177–183.

46. Kühner, F., Costa, L.T., Bisch, P.M., Thalhammer, S., Heckl, W.M. and Gaub, H.E. (2004) LexA-DNA bond strength by single molecule force spectroscopy. Biophys. J., 87, 2683–2690.

47. Ros, R., Eckel, R., Bartels, F., Sischka, A., Baumgarth, B., Wilking, S.D., Pühler, A., Sewald, N., Becker, A. and Anselmetti, D. (2004) Single molecule force spectroscopy on ligand-DNA complexes: From molecular binding mechanisms to biosensor applications. J. Biotechnol., 112, 5–12.

48. Chalmers, R.M. and Kleckner, N. (1994) Tn10/IS10 transposase purification, activation, and in vitro reaction. J. Biol. Chem., 269, 8029–8035.

49. Mizuuchi, K. (1992) Transpositional recombination: Mechanistic insights from studies of Mu and other elements. Annu. Rev. Biochem., 61, 1011–1051.

50. Harshey, R.M. (2014) Transposable phage Mu Rasika. Microbiol Spectr, 2.

51. Fuller, J.R. and Rice, P.A. (2017) Target DNA bending by the Mu transpososome promotes careful transposition and prevents its reversal. Elife, 6, e21777.

52. Bouuaert, C.C., Lipkow, K., Andrews, S.S., Liu, D. and Chalmers, R. (2013) The autoregulation of a eukaryotic DNA transposon. Elife, 10.7554/eLife.00668.

