## Supplementary file for "AFM-based force spectroscopy unravels the stepwise-formation of a DNA transposition complex driving multi-drug resistance dissemination"

Fernandez et al.

**This PDF file includes:**

Figures S1 to S9

Tables S1 to S4

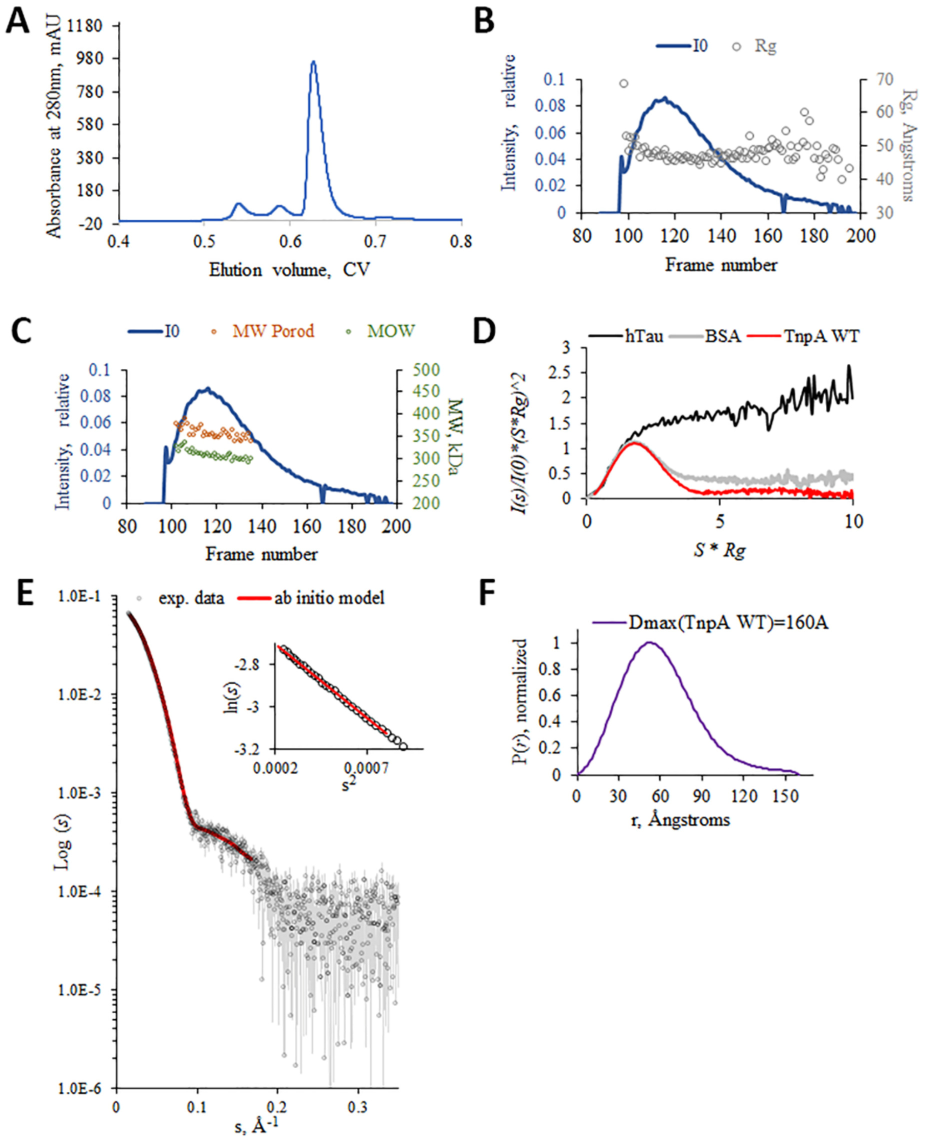


**Fig. S1 | Analysis of SEC-SAXS data for TnpA$^{WT}$.** (A) The absorbance at 280nm as a function of elution volume. (B and C) The X-ray scattering intensity as a function of elution time (exposure frame) from the size exclusion and radius of gyration (B) or molecular weight estimation (C) using Porod volume and apparent volume (MoW). The dashes purple lines indicate the region used for frame averaging. (D) Dimensionless Kratky plot overlay of TnpA$^{WT}$ compared to references: the globular BSA (grey line) and highly flexible hTau40wt proteins (black line)(1). (E) SAXS profile for TnpA$^{WT}$ (experimental data, gray circles; experimental errors, thin gray lines) and the fit to the best *ab initio* model with a $\chi$ of 1.63 (Fig. 1*B*). Inset shows a linear Guinier plot confirming absence of interparticle interactions. (F) The pair-distance distribution function, P(r), with respective Dmax value.

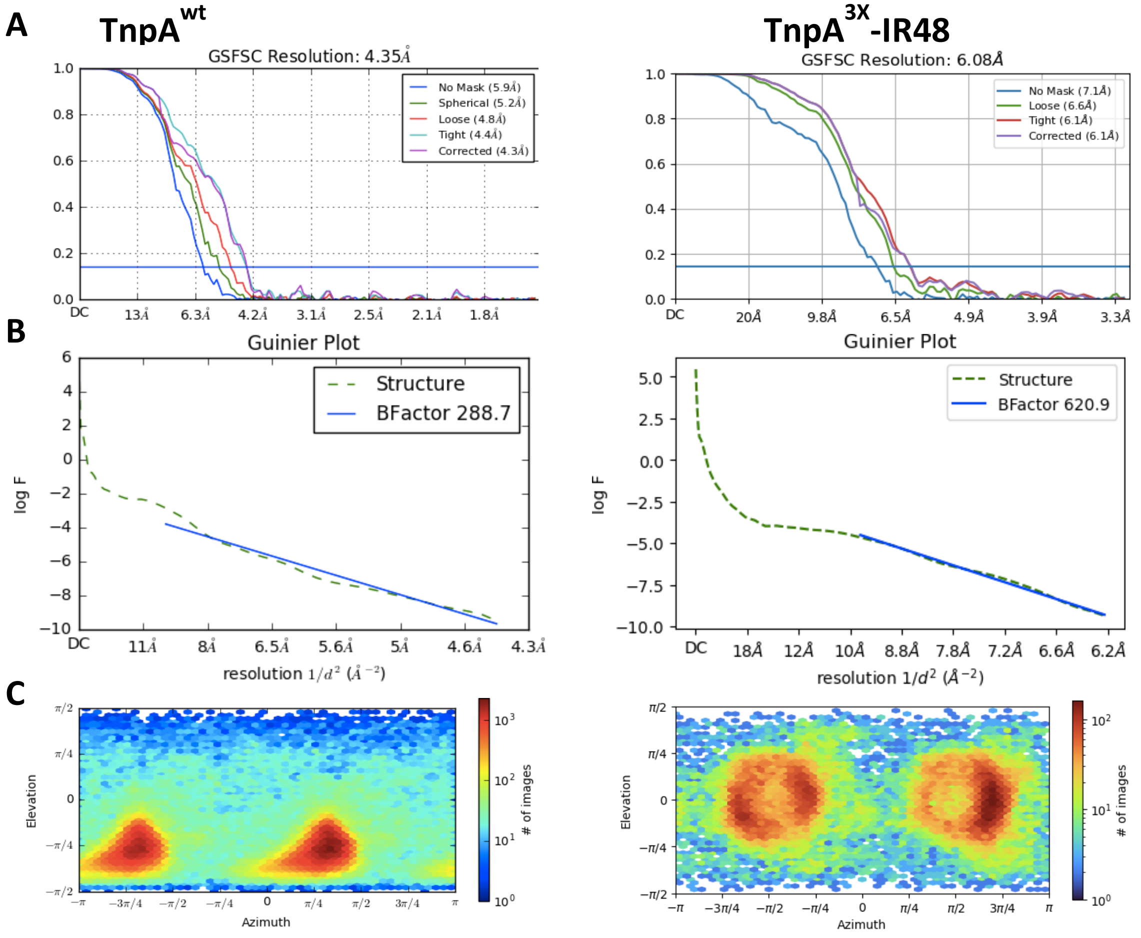


**Fig. S2 | Cryo-EM processing results for TnpA$^{WT}$ (left) and TnpA$^{3X}$-IR48 (right).** (A) FSC curves, (B) Guinier plots and calculated B factors, (C) Angular distribution or particle orientations as calculated in cryoSPARC.

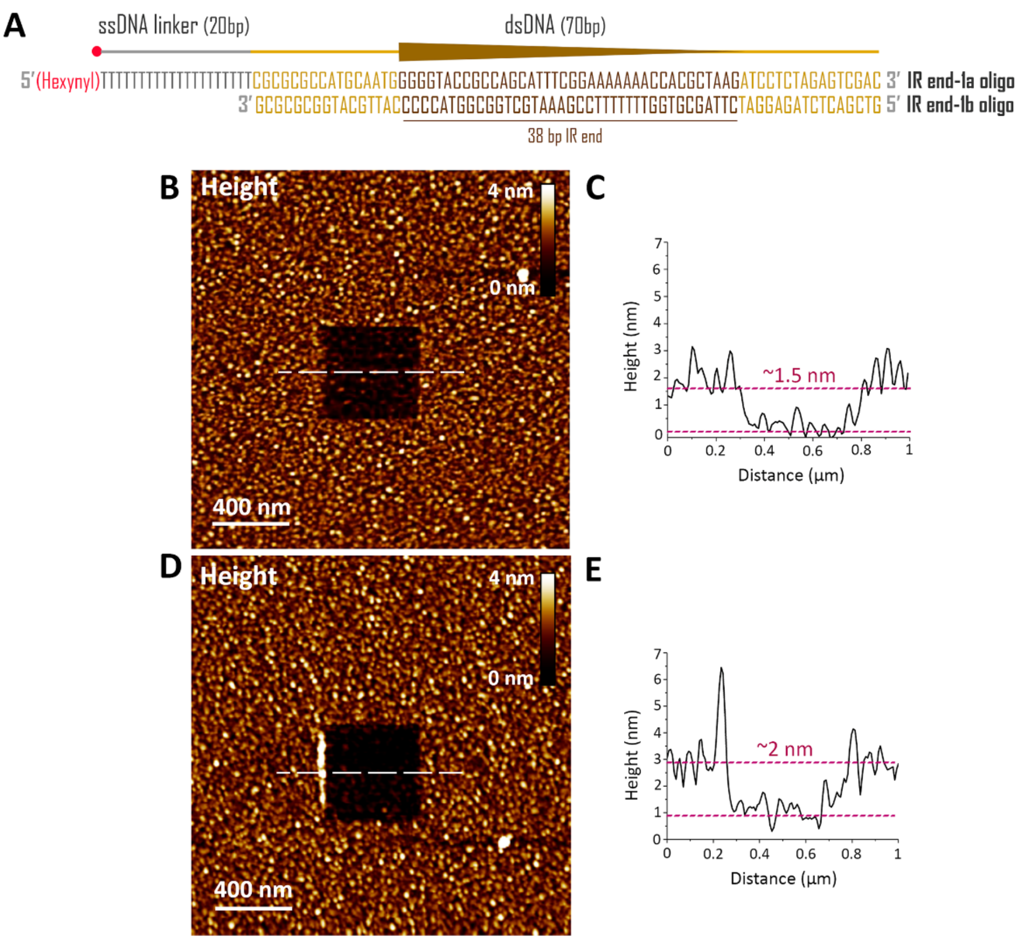


**Fig. S3 | Validation of the one-end substrate immobilization on the surface.** Specific DNA substrate containing a Tn4430 IR end was assembled by annealing two complementary oligonucleotides (A and Table S4). The substrate carries an alkyne reactive group (Hexynyl) at their 5' end after a 20-nt single-stranded DNA linker. AFM topography image of a PEG/PEG-$N_3$ (B) or PEG/PEG-$N_3$/one end (D) coated surface after scanning a 0.5 x 0.5 µm area at high forces to remove the attached biomolecules (referred to as 'scratching experiment'). The biomolecule-free surface of inside the square was ~1.5 nm (C) or ~2 nm (E) lower than the surrounding biomolecule-coated surface, which provides an estimated thickness for the deposited PEG and one-end substrate, respectively.

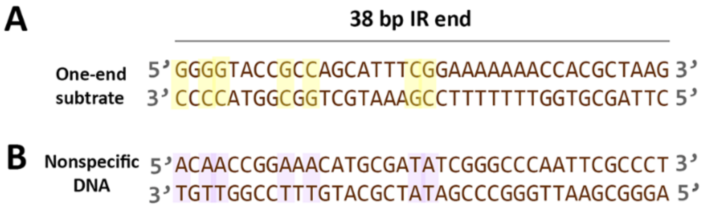


**Fig. S4 | Nonspecific DNA design.** (A) The IR sequence of the one-end substrate with the conserved motifs highlighted in yellow. (B) In the nonspecific DNA, the seven conserved positions were replaced (i.e. Gs by As and Cs by Ts; they shown in purple), also the GC/AT balance was equilibrated of the nonspecific oligo compared to the specific substrate.

**DNA molecules containing a second transposon end block SEC formation**

To determine whether linear DNA molecules containing a second transposon end can interfere with SEC formation by blocking binding sites on TnpA, a second DNA molecule was designed and synthesized. The molecule, called IR end-2 or blocking DNA (sequence provided in Table S4), mimicked the second end of the transposon. In this way, blocking DNA was used to study the binding inhibition properties through our single-molecule force spectroscopy approach. First, the binding probability (BP) between TnpA and the one-end was measured in the absence of the blocking DNA (0 μM), and then the latter was injected at three different concentrations (12.5, 25, and 50 μM) (Fig. S5A). For the three TnpA proteins (WT, 3X and S911R), a progressive BP reduction was observed as a function of the concentration in presence of the blocking DNA (Fig. S5B-D), which confirms a specific inhibition.

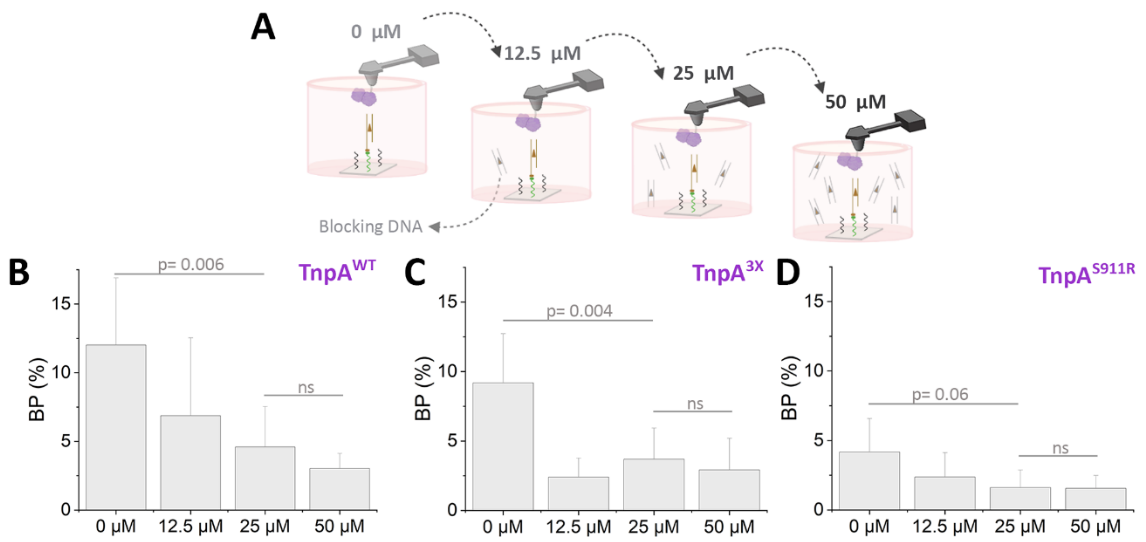


**Fig. S5 | Monitoring the effect of the addition of the second transposon end on SEC formation.** (A) Efficiency of blocking DNA (IR end-2) is evaluated by measuring the binding probability of the TnpA-one end interaction before and after the incubation of the functionalized AFM tip with IR end-2 at increasing concentrations (12.5, 25 or 50 µM). (B-D) Histograms showing binding probability (BP) without the blocking DNA (0 µM) and upon incubation with 12.5, 25 or 50 µM of blocking DNA for $TnpA^{WT}$ (B), $TnpA^{3X}$ (C) and $TnpA^{S911R}$ (D). The error bar indicates s.d. of the mean value. Data are representative of at least n= 3 independent experiments (tips and samples) per DNA concentration. P values were determined by two-sample t test with R software.

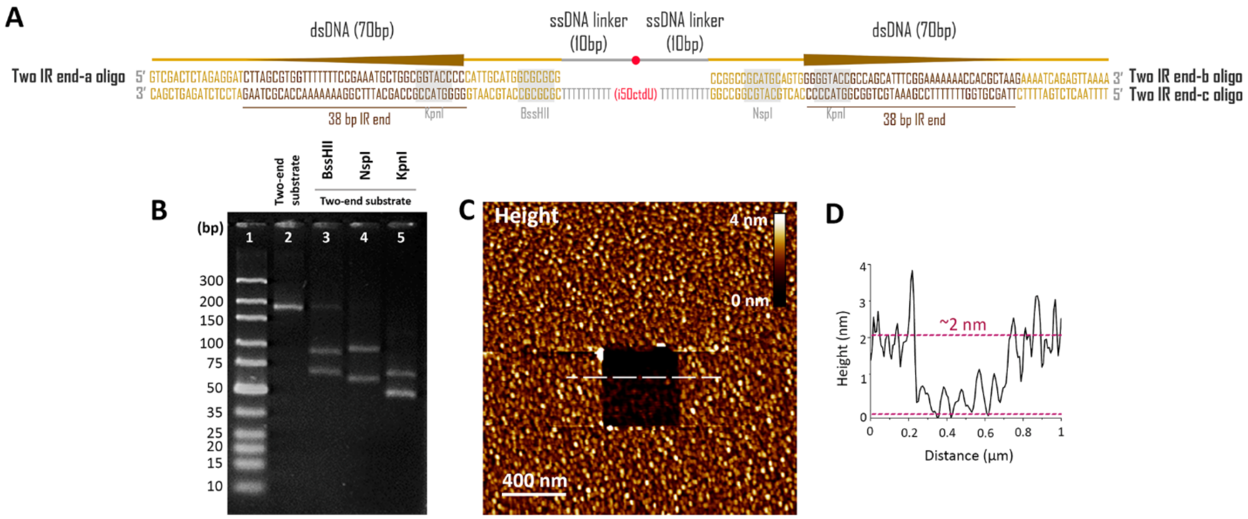


**Fig. S6 | Oligos annealing control and validation of the immobilization of the two-end substrate on the surface.** (A) Specific DNA substrate containing properly oriented Tn4430 IR ends was assembled by annealing three complementary oligonucleotides (Table S4). The duplex regions corresponding to the IR ends are separated by 20-nt single-stranded DNA linker, which contain an internal alkyne group (i5OctdU). (B) Gel electrophoresis of the ~160 bp substrate containing two transposon ends. Five percent agarose gel of the digested and undigested DNA substrate. Lane 1, = markers are low molecular weight DNA ladder (NEB); lane 2, undigested two ends substrate; Lanes 3-5, digested two ends substrate with BssHII, NspI and KpnI (NEB), respectively. The gel was stained with 0.5 µg/mL ethidium bromide (Sigma, Poole, UK) and run at 120 volts for 40 minutes. (C) AFM topography image of a PEG/PEG-N3/two ends coated surface after scanning a 0.5 µm x 0.5 µm area at high forces to remove the attached biomolecules. (D) Cross-section taken along the white dashed line in C. The biomolecule-free surface inside the square was ~2 nm lower in height than the surrounding biomolecule-coated surface, which provides an estimated thickness for the deposited two ends layer.

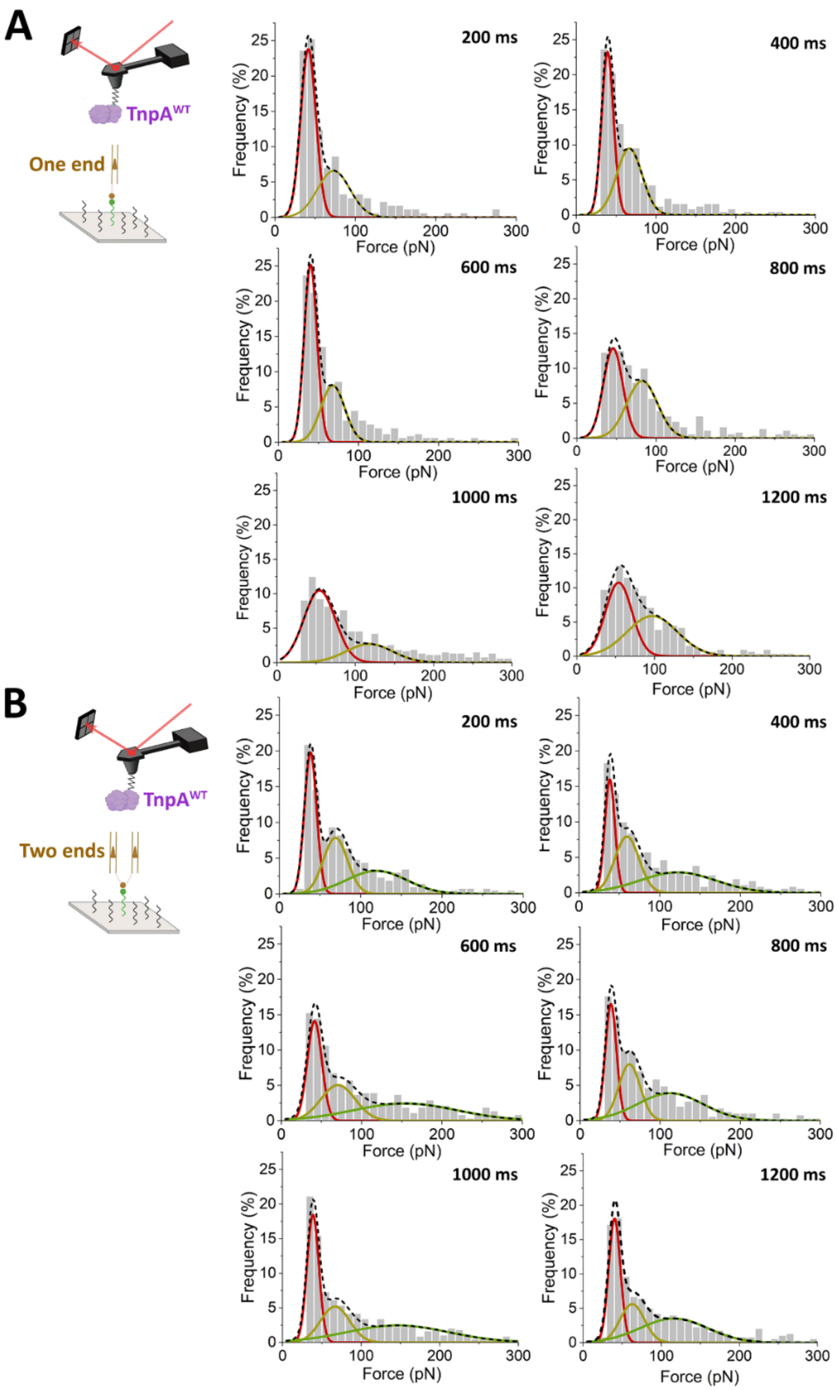


**Fig. S7 | Comparison of binding frequency between TnpA$^{WT}$ and the one-end substrate (A) or the two-end substrate (B).** Frequency distributions of rupture forces at contact times of 200 ms, 400 ms, 600 ms, 800 ms, 1000 ms and 1200 ms.

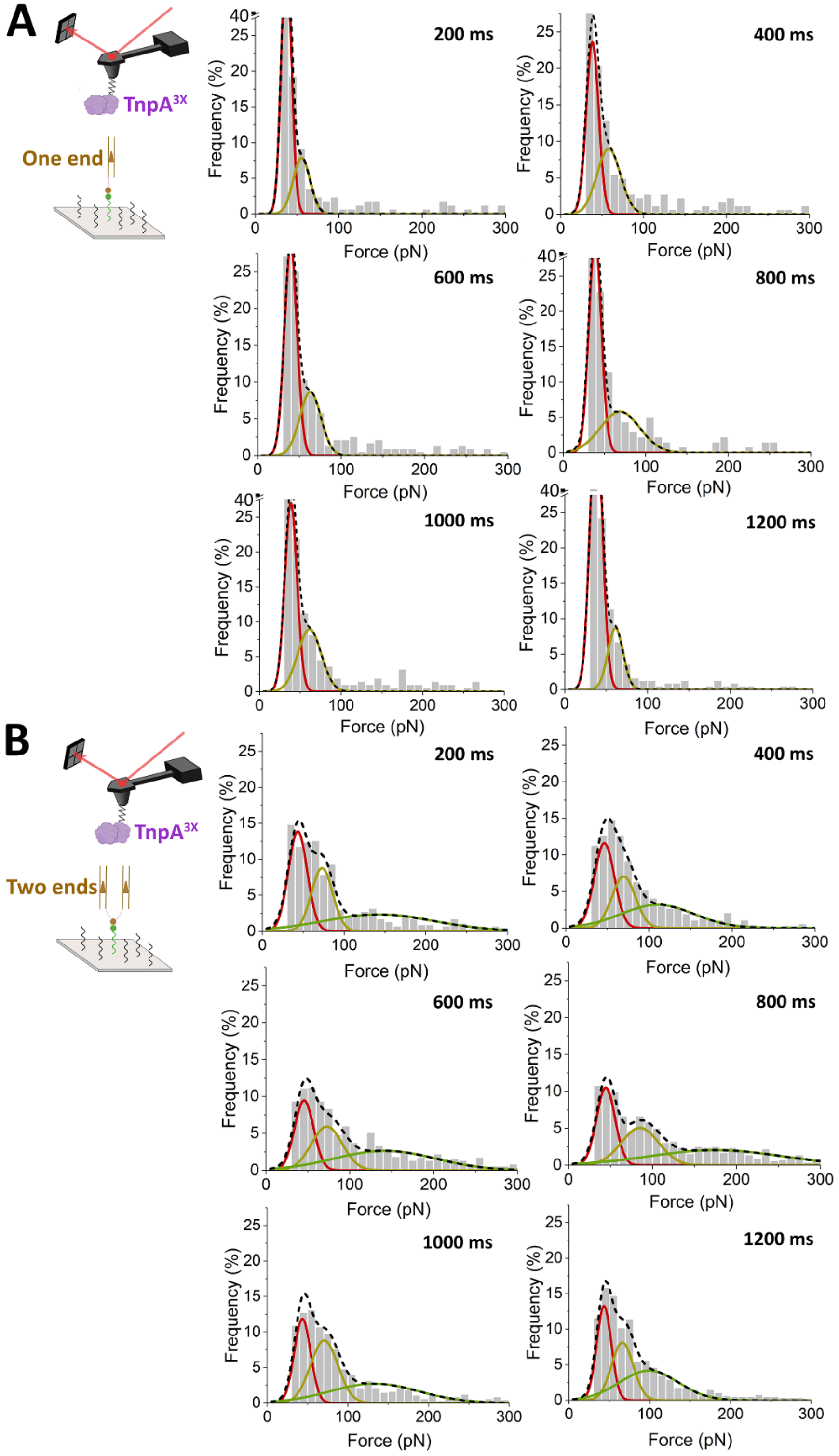


**Fig. S8 | Comparison of binding frequency between TnpA$^{3X}$ and the one-end substrate (A) or the two-end substrate (B).** Frequency distributions of rupture forces at contact times of 200 ms, 400 ms, 600 ms, 800 ms, 1000 ms and 1200 ms.

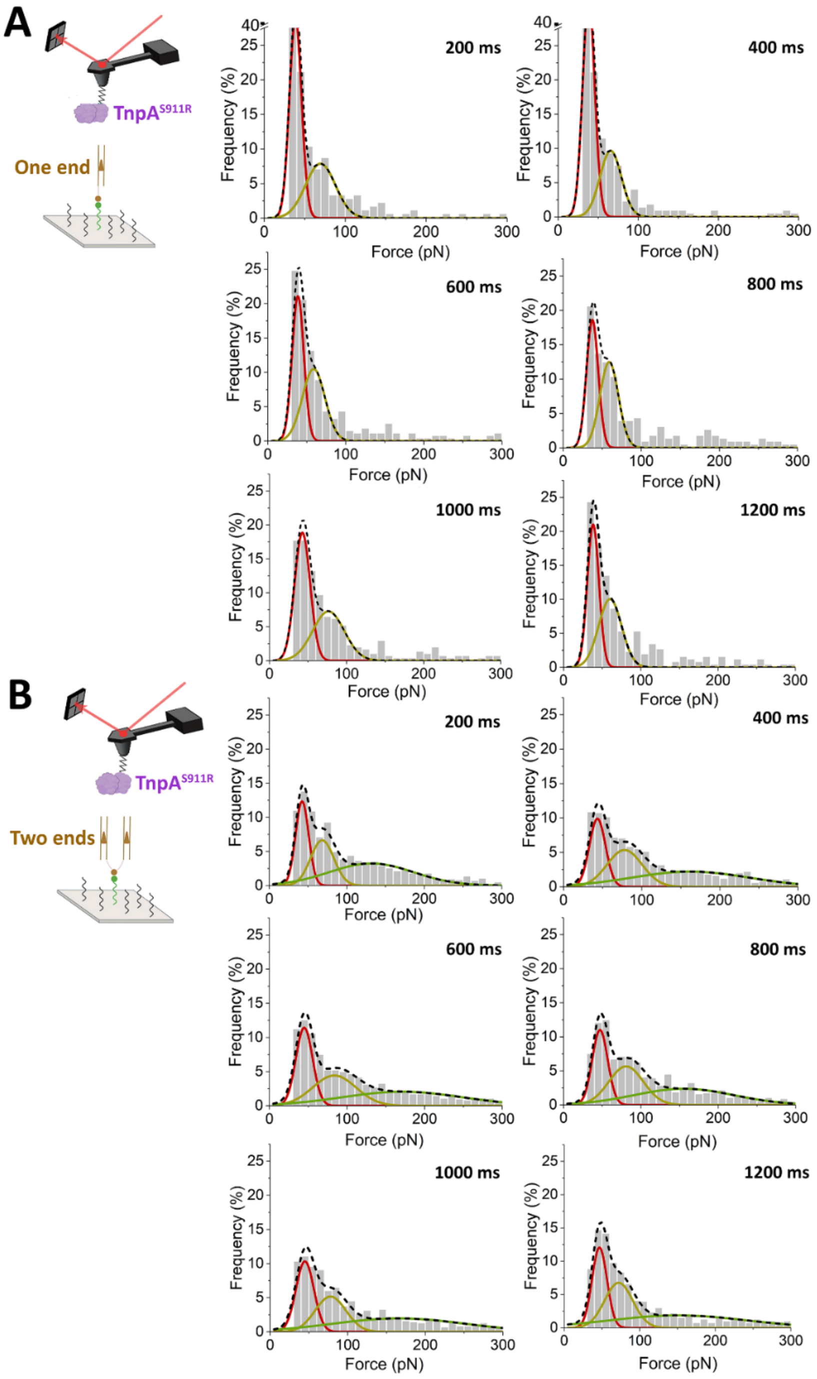


**Fig. S9 | Comparison of binding frequency between TnpA$^{S911R}$ and the one-end substrate (A) or the two-end substrate (B).** Frequency distributions of rupture forces at contact times of 200 ms, 400 ms, 600 ms, 800 ms, 1000 ms and 1200 ms.

**Table S1. Overall parameters and modeling summary of SAXS data for TnpA$^{WT}$**

| Data-collection parameters | TnpA$^{WT}$ |
|---|---|
| Instrument: | Swing, Soleil |
| Wavelength (Å) | 1.022 |
| *s* range (Å$^{-1}$)* | 0.0005-0.6093 |
| Temperature (K) | 288 |
| Detector: | EigerX4M |
| Concentration (mg/ml)/Vol. (ul): | 3.6/50 |
| *Structural parameters:* | |
| *I(0)* (Å$^{-1}$) [from *P(r)*]: | 0.07 ± 0.01 |
| *Rg* (Å) [from *P(r)*]: | 46.09 ± 0.09 |
| *I(0)* (Å$^{-1}$) (from Guinier): | 0.07 ± 0.01 |
| *Rg* (Å ) (from Guinier): | 45.81 ± 0.08 |
| $D_{max}$ (Å): | 160.00 ± 10 |
| Porod volume estimate, $V_p$ (Å$^3$): | 480,000 ± 20,000 |
| Excluded volume, $V_{ex}$ (Å$^3$): | 540,000 ± 20,000 |
| *Molecular-mass determination:* | |
| Molecular mass $M_r$ (Da) [from SaxsMOW]: | 300,000 ± 10,000 |
| Molecular mass $M_r$ (Da) from Porod volume ($V_p$/1.8) | 270,000 ± 10,000 |
| Molecular mass $M_r$ (Da) from excl. volume ($V_{ex}$/1.8) | 300,000 ± 10,000 |
| Calculated $M_r$ (kDa) from sequence: | 116.76 |
| *Modeling parameters:* | |
| Shape reconstruction | GASBOR |
| Symmetry | P2 |
| range of $\chi$ | 1.63-2.17 |
| # of models averaged/total | 10/10 |
| DAMAVER NSD (var) | 1.12 (0.02) |
| *Software used:* | |
| Primary data reduction: | FOXTROT |
| Data processing and evaluation: | DATSW, PRIMUS, GENOM |
| Shape reconstruction: | DAMMIN, GASBOR |
| 3D graphics: | PYMOR |
| *Small Angle Scattering Biological Data Bank* | |
| SASBDB accession codes | SASDMR5 |

Abbreviations: Vol., injected volume *I(0)*, extrapolated scattering intensity at zero angle; *Rg*, radius of gyration calculated using either Guinier approximation (from Guinier) or the indirect Fourier transform package GNOM [from P(r)]; $M_r$, molecular mass; *Dmax*, maximal particle dimension; $V_p$, Porod volume; $V_{ex}$, particle excluded volume.
*Momentum transfer $|s| = 4\pi\sin(\theta)/\lambda$.

**Table S2. Statistics of cryo-EM data collection.**

| | $TnpA^{WT}$ | $TnpA^{S911R}$ | $TnpA^{S911R}$ IR48 | $TnpA^{3X}$ IR48 |
|---|---|---|---|---|
| **Data collection** | | | | |
| Microscope | JEOL CRYOARM300 | | | |
| Energy filter / slit width [eV] | In-column Omega energy filter / 20 | | | |
| Detector | K2 summit (Gatan) | K3 | | |
| Nominal magnification | 60 000 | 60 000 | | |
| Accelerating voltage [kV] | 300 | 300 | | |
| Calibrated pixel size [Å] | 0.786 | 0.764 | 0.784 | 0.784 |
| Defocus range [µm] | 1-3.5 | 1-4 | 1.4-2.6 | 0.8-4 |
| Frames per movie | 50 | 61 | 50 | 61 |
| Exposure time, *s* | 10 | 2.985 | 2.985 | 2.985 |
| Total electron dose [$e^-/Å^2$] | 58 | 64.6 | 64.6 | 64.6 |
| Automation software (SerialEM) | v3.7.6 | v3.8.0 | | |
| Total movies used | 4,629 | 1,700 | 6,429 | 4,155 |
| **Reconstruction** | | | | |
| Final particles [no.] | 267,639 | - | - | 45,955 |
| Box size [px] | 320 | - | - | 500 |
| Imposed symmetry | C2 | - | - | C2 |
| Sharpening B-factor, cryoSPARC [$Å^2$] | -289 | - | - | -621 |
| Resolution, cryoSPARC [Å] (GFSC=0.143) | 4.4 | - | - | 6.1 |

### Table S3. Bacterial strain and plasmids used in this study

| Name | Description | Reference o source |
|---|---|---|
| E. coli strain TOP10 | F- mcrA Δ(mrr-hsdRMS-mcrBC) φ80lacZM15 ΔlacX74 nupG araD139 Δ(ara-leu)7697 galE15 galK16 rpsL endA1 λ- | Invitrogen |
| pGIAD003 | $Tc^r$, expression of wild-type TnpA-myc-$His_6$ under p*Ara* promoter | (2) |
| pGIMLS911R/M-H | $Tc^r$, expression of mutant $TnpA^{S911R}$-myc-$His_6$ under p*Ara* promoter | (3) |
| pGIML3×/M-H | $Tc^r$, expression of mutant $TnpA^{3\times}$(W24R-A174V-S911R)-myc-$His_6$ under p*Ara* promoter | (3) |

$Tc^r$, resistance to tetracycline.

### Table S4. Oligonucleotides used in this study

| Name | 5' to 3' sequence | Description |
|---|---|---|
| *IR end-1a* | (Hexynyl)TTTTTTTTTTTTTTTTTTTTCGCGCGCCATGCAATGGGGGTACCGCCAGCATTTCGGA AAAAAACCACGCTAAGATCCTCTAGAGTCGAC | Annealing with *IR end-1b* generates a 70-bp IR-containing DNA substrate |
| *IR end-1b* | GTCGACTCTAGAGGATCTTAGCGTGGTTTTTTTCCGAAATGCTGGCGGTACCCCCATTGCATGGCGCGCG | Annealing with *IR end-1a* generates a 70-bp IR-containing DNA substrate |
| *IR end-2a* | CCGGCCGCATGCAGTGGGGGTACCGCCAGCATTTCGGAAAAAAACCACGCTAAGAAAATCAGAGTTAAAA | Annealing with *IR end-2b* generates a 70-bp IR-containing DNA substrate |
| *IR end-2b* | TTTTAACTCTGATTTTCTTAGCGTGGTTTTTTTCCGAAATGCTGGCGGTACCCCCACTGCATGCGGCCGG | Annealing with *IR end-2a* generates a 70-bp IR-containing DNA substrate |
| *Nonspecific DNA-a* | (Hexynyl)TTTTTTTTTTTTTTTTTTTTCGCGCGCCATGCAATGACAACCGGAAACATGCGATATCGGGCCCAATTCGCCCTATCCTCTAGAGTCGAC | Annealing with *nonspecific DNA-a* generates a 70-bp DNA nonspecific substrate |
| *Nonspecific DNA-b* | GTCGACTCTAGAGGATAGGGCGAATTGGGCCCGATATCGCATGTTTCCGGTTGTCATTGCATGGCGCGCG | Annealing with *nonspecific DNA-b* generates a 70-bp DNA nonspecific substrate |
| *Two IR end-a* | GTCGACTCTAGAGGATCTTAGCGTGGTTTTTTTCCGAAATGCTGGCGGTACCCCCATTGCATGGCGCGCG | Annealing with *two IR end-b* and *c* generates a 160-bp IR double-containing DNA substrate |
| *Two IR end-b* | CCGGCCGCATGCAGTGGGGGTACCGCCAGCATTTCGGAAAAAAACCACGCTAAGAAAATCAGAGTTAAAA | Annealing with *two IR end-a* and *c* generates a 160-bp IR double-containing DNA substrate |
| *Two IR end-c* | TTTTAACTCTGATTTTCTTAGCGTGGTTTTTTTCCGAAATGCTGGCGGTACCCCCACTGCATGCGGCCGGTTTTTTTTTT(i5OctdU)TTTTTTTTTCGCGCGCCATGCAATGGGGGTACCGCCAGCATTTCGGAAAAAAACCACGCTAAGATCCTCTAGAGTCGAC | Annealing with *two IR end-a* and *b* generates a 160-bp IR double-containing DNA substrate |

(Hexynyl): 5' terminal alkyne group; (i5OctdU): internal alkyne group.

Oligonucleotides were purchased from Eurogentec (Seraing, Belgium). End-modifications are highlighted in red.

### REFERENCES

1. Gorantla,N.V., Shkumatov,A. V. and Chinnathambi,S. (2017) Conformational dynamics of intracellular tau protein revealed by CD and SAXS. *Methods Mol. Biol.*, **1523**, 3–20.
2. Nicolas,E., Oger,C.A., Nguyen,N., Lambin,M., Draime,A., Leterme,S.C., Chandler,M. and Hallet,B.F.J. (2017) Unlocking Tn3-family transposase activity in vitro unveils an asymetric pathway for transposome assembly. *Proc. Natl. Acad. Sci. U. S. A.*, **114**, 669–678.
3. Lambin,M., Nicolas,E., Oger,C.A., Nguyen,N., Prozzi,D. and Hallet,B. (2012) Separate structural and functional domains of Tn4430 transposase contribute to target immunity. *Mol. Microbiol.*, **83**, 805–820.
